# Short term memory properties of sensory neural architectures

**DOI:** 10.1101/590851

**Authors:** Alexis M. Dubreuil

## Abstract

A functional role of the cerebral cortex is to form and hold representations of the sensory world for behavioral purposes. This is achieved by a sheet of neurons, organized in modules called cortical columns, that receives inputs in a peculiar manner, with only a few neurons driven by sensory inputs through thalamic projections, and a vast majority of neurons receiving mainly cortical inputs. How should cortical modules be organized, with respect to sensory inputs, in order for the cortex to efficiently hold sensory representations in memory? To address this question we investigate the memory performance of trees of recurrent networks (TRN) that are composed of recurrent networks, modeling cortical columns, connected with each others through a tree-shaped feed-forward backbone of connections, with sensory stimuli injected at the root of the tree. On these sensory architectures two types of short-term memory (STM) mechanisms can be implemented, STM via transient dynamics on the feed-forward tree, and STM via reverberating activity on the recurrent connectivity inside modules. We derive equations describing the dynamics of such networks, which allow us to thoroughly explore the space of possible architectures and quantify their memory performance. By varying the divergence ratio of the tree, we show that serial architectures, where sensory inputs are successively processed in different modules, are better suited to implement STM via transient dynamics, while parallel architectures, where sensory inputs are simultaneously processed by all modules, are better suited to implement STM via reverberating dynamics.

## 1 Introduction

Neural activity in the neocortex forms sensory representations of ongoing stimuli that can be held in memory for behavioral purposes (Fuster, 1995). Such operations are performed by cortical circuits which possess a peculiar organization, as described by anatomical studies. For instance cortical connectivity has been shown to be spatially clustered, which allows to conceptually segregate the cortical sheet into groups of neurons called cortical columns (Bosking et al., 1997; DeFelipe et al., 1986; Gilbert and Wiesel, 1989; Pucak et al., 1996). Regarding the transmission of sensory information to cortical circuits, anatomical studies have suggested that a small subset of neurons are directly driven by thalamic projections, while a vast majority are in charge of processing these stimuli (Braitenberg and Schütz, 1991). This vast majority of neurons are sometimes called interneurons as they are not directly under the influence of sensory inputs, nor directly involved in generating a cortical output. In this paper we investigate how the organization of these interneurons, with respect to sensory inputs, impact the memory performance of a cognitive architecture. Two extreme types of organization can be considered. A serial organization, where a sensory input is successively passed to different groups of neurons, with one group operating on the output of the previous one (Fig. 1a). A parallel organization, where a sensory input is simultaneously sent to all groups of neurons, where it is processed independently from one group to another (Fig. 1b). The neural architectures we consider range from serial to parallel, and consist of modules, made out of recurrent neural networks modeling cortical columns, that for simplicity we consider connected in a feed-forward manner. These neural architectures implement two kinds of short-term memory mechanisms. First, the recurrent connectivity inside each module creates feed-back loops in which neural activity can reverberate leading to sustained patterns of activity coding for specific stimuli, i.e. modules are attractor neural networks (Amit, 1989; Amit and Brunel, 1997; Hopfield, 1982; Tsodyks and Feigel*’*man, 1988). Second, the feed-forward connectivity between modules allows to implement delay-lines on which sequences of stimuli propagate. Delay-lines have been shown to be efficient neural architectures to hold sensory representations in the transient dynamics of a neural network (Ganguli et al., 2008; Jaeger and Haas, 2004; Lim and Goldman, 2012; Maass et al., 2002; White et al., 2004). The advantage of the attractor neural network scenario is that memory can be held on a very long timescale. A drawback is that only a limited number of already learnt sensory representations can be maintained, since maintenance of a representation relies on a specific set of synaptic connections. We will refer to this number as the *storage capacity* of the architecture. On the opposite, short-term memory via transient dynamics allows to memorize arbitrary stimuli, but the timescale on which these stimuli can be maintained is limited. We will refer to this time scale as the *buffering capacity* of the network.

**Figure 1.**
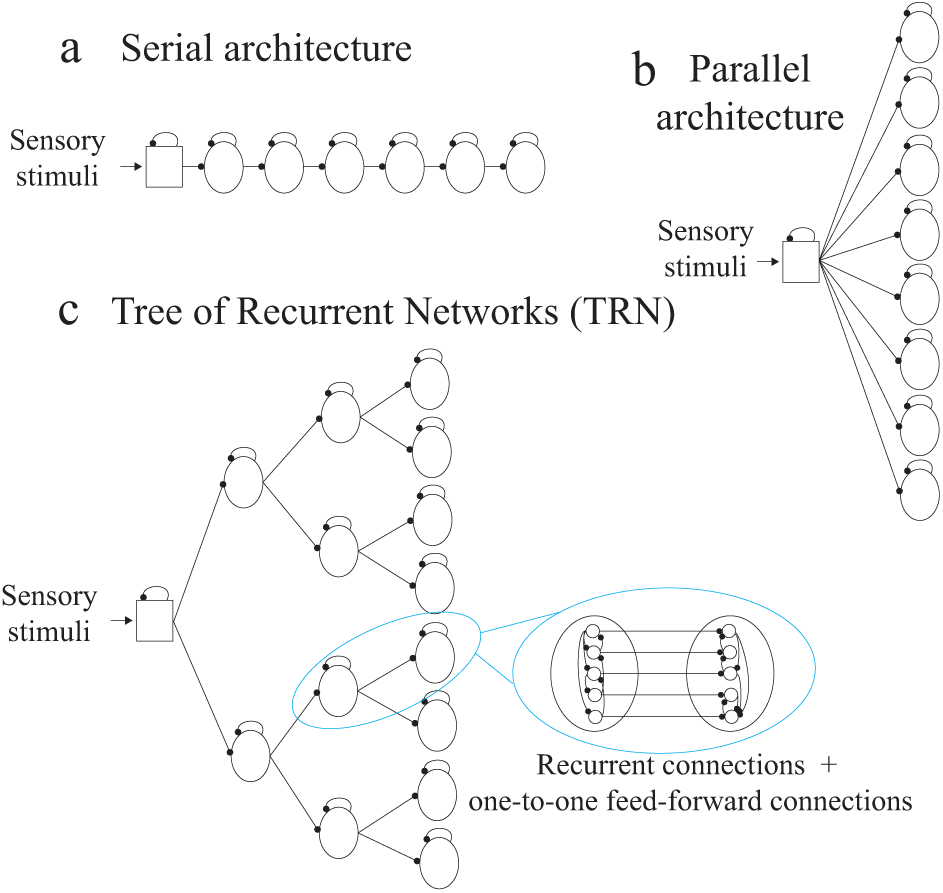
Trees of recurrent networks (TRN) **a** Purely serial architecture, obtained as a special case of TRN with divergence ratio d = 1. The root module (squared box), models the thalamus conveying sensory inputs to modules modeling cortical columns. Sensory inputs propagate from left to right through the feed-forward connections between modules. **b** Purely parallel architecture, obtained as a special case of TRN with divergence ratio d = M, where M is the number of modules composing the network. **c** TRN with divergence ratio d = 2, M = 14 and L = 3. Each module is an attractor network of N neurons with recurrent connectivity storing αN patterns of activity as fixed points of the module’s dynamics. Connections between a module and each of its d children are one-to-one, i.e. a neuron of the parent module projects to a single neurons in each child module and each neuron of a child module receives a single projection.

In the first section of this paper we formally define the class of neural architectures called Tree of Recurrent Networks (TRN) that we study. In order to reach a full comprehension of the memory properties of these architectures we take the following incremental steps. We first study the storage capacity of an isolated module, as well as the dynamics of memory retrieval (second section). We then focus on a single feed-forward path of a TRN, showing in particular how the recurrent connections inside modules degrade stimuli as they propagate throughout the path. An important phenomenon that eventually limits the storage and buffering capacity of the path (third section). We finally examine how these two quantities behave in a full TRN, in particular we compare the memory properties of the extreme cases that are the fully serial and fully parallel architectures (fourth section). To conclude we summarize our results and discuss how TRN can be mapped onto cortical sensory hierarchies.

## 2 Definition of the model

We consider feed-forward trees of *M* modules (Fig. 1) such that for each step up the tree, the number of modules increases by a factor *d* where *d* labels the divergence ratio of the TRN. The divergence ratio ranges from *d* = 1, defining a purely serial architecture composed of a single path of length *L* = *M* (Fig. 1a), to *d* = *M*, defining a purely parallel architecture composed of *M* paths of length *L* = 1(Fig. 1b). For intermediate divergence ratios, the length of the paths is related to *M* and d as detailed in section 5, and illustrated in Fig. 1c.

Each module operates as a fixed-points attractor neural network in which independent memories are stored in the recurrent connections 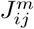(*m* = 1, …, *M* and *i,j* = 1, …,*N*, where *N* is the number of neurons in each module).

### 2.1 Dynamics of the network

Neuron *i* in module m is described by a binary variable 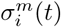 whose dynamics is:

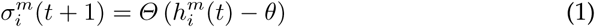

where *Θ*(.) is the Heaviside function, *θ* an activation threshold and 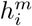 is the total synaptic input on the neuron, which is the sum of a feed-forward input, coming from a single neuron of the previous module, and a recurrent input:

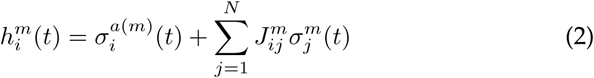

where module *a*(*m*) is the module preceding *m* in the feed-forward structure.

### 2.2 Recurrent connections

The recurrent connections of a module store *P* random independent binary patterns, {***ξ***^*μ,m*^}, *μ* = 1, …, *P* with a coding level *f*,

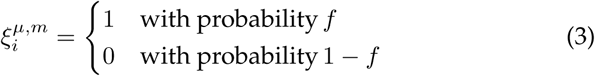

Patterns are stored using a covariance learning rule (Sejnowski, 1977; Tsodyks and Feigel*’*man, 1988) which implements feed-back loops allowing activity to reverberate such that the *P* patterns can become fixed points of the network dynamics

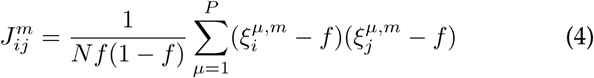

We introduce the storage load *α* = *P/N* which quantifies the number of memories stored in each module. Note that in what follows, the coding level is set to *f* = 0.01 unless stated otherwise, we have checked that other choices of coding levels (*f* = 0.001 or *f* = 0.1) lead to qualitatively similar results.

### 2.3 Inputs to the network

At each time step, a new input is represented by the state of the root module ***σ***^0^. In order to probe short-term memory properties via transient dynamics and reverberating activity, we consider the TRN under two kinds of stimulation. In the first scenario, inputs are drawn independently of the stored patterns (see Fig. 4a). In the second scenario, the network receives a similar stream of inputs, but at *t* = 0 the root module represents a pattern stored in module *m*_0_ (see Fig. 5a).

## 3 Storage capacity of a single module

The recurrent connectivity of each module is the substrate allowing TRN to hold stimuli in short-term memory via reverberating activity. With our choice of connectivity matrix (4), modules operate as attractor neural networks and αN stimuli can be held in short-term memory under the form of persistent activity. We quantify the ability of a module to implement such short-term memory mechanism with the storage capacity (Amit, 1989). It is defined as the maximal storage load *α* such that imprinted patterns are stable states of the network’s dynamics. We also describe the basins of attraction associated to each stored memory.

### 3.1 Dynamical equations for macroscopic variables

Without loss of generality we focus on the retrieval and maintenance of a specific memory, e.g. pattern ***ξ***^1^. The state of the module is described by the overlap *m*(*t*) between the instantaneous network state and the corresponding pattern

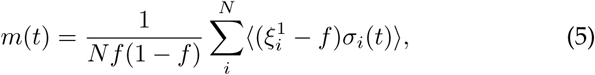

and the average activity in the module,

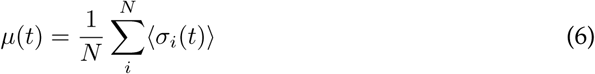

where 〈.〉 corresponds to averaging over different microscopic initial states (that have the same macroscopic description). The dynamics of these order parameters can be derived using standard methods (see Supplementary materials, (Evans, 1989; Sompolinsky and White, 2003)):

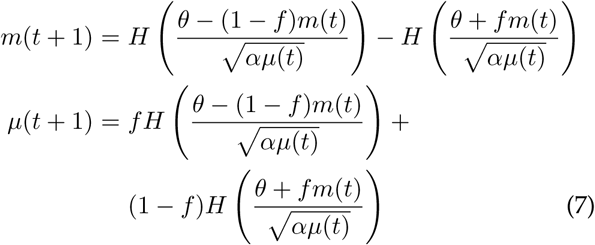

where

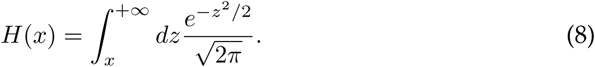

### 3.2 Stable states and storage capacity

We label (*m*\*, *μ*\*) fixed point of (7) for the overlap and mean activity. A particular fixed point is the background state *m*\* = 0 and *μ*\* = 0. Retrieval states that have a high overlap with a stored pattern *m*\* ≃ 1 exist as long as the storage load remains below a critical value *α* < *α*_*c*_. This critical value is shown by the blue curve in Fig. 2a that we obtain by numerically finding the fixed points of equation (7) for *f* = 0.01. The retrieval states have been described previously for *f* ≪ 1 (Tsodyks and Feigel*’*man, 1988; White et al., 2004). In this regime the storage capacity *α_c_* can be approximated by

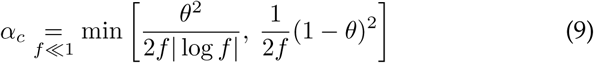

**Figure 2.**
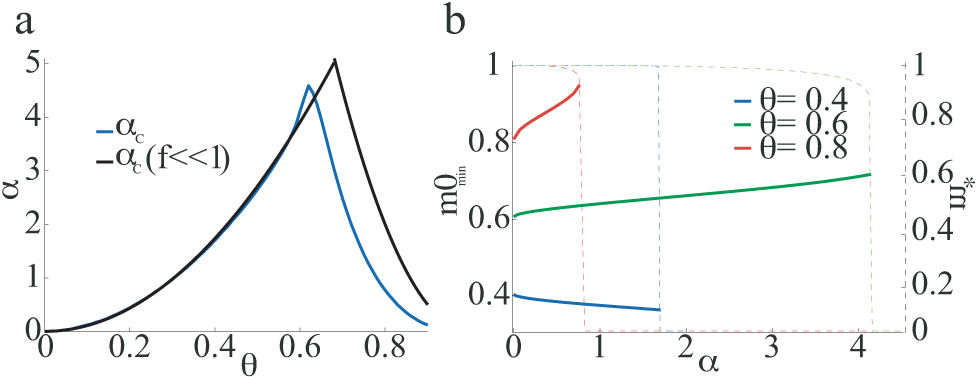
Short-term memory via reverberating activity in a single module **a** The blue curve shows the storage capacity for f = 0.01 obtained by iterating (7). The black curve shows the analytical approximation (9) obtained for f ≪ 1. **b** Size of the basins of attraction associated to each stored patterns (full lines), together with the overlap m* between the fixed point of the network and the memory pattern to be stored (dashed lines). We measure the size of the basins of attraction as the minimal initial overlap m0_*min*_ that leads to pattern retrieval when solving (7) with μ(t = 0) = f.

The curve *α*_*c*_(*f* ≪ 1) vs *θ* is shown in black in Fig. 2a. It is composed of two quadratic branches in *θ*. On the first one, retrieval states are destabilized by an increase in the activity of background neurons (neurons *i* such that 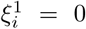), while on the second branch, retrieval states are destabilized by a silencing of foreground neurons (neurons *i* such that 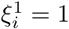).

By solving equations (7) numerically, we have also found an *’*active*’* background state (*m*\* = 0, *μ*\* ≫ *f*) and *’*weak*’* retrieval states characterized by *(m*\* = *O*(1),*μ*\* ≫ *f*). We study such states in Supplementary materials. Note that these states exist at low activation thresholds, which are sub-optimal for TRN memory properties, as will be shown in the following.

### 3.3 Basins of attraction

The state in which the network settles depends on the two initial values *m*_0_ *= m(t* = 0) and *μ*_0_ = *μ*(*t* = 0). We measure the size of the basins of attraction by *m*0_*min*_ the minimal value of the overlap that is required to reach a retrieval state for an initial mean activity *μ*_0_ = *f*. In Fig. 2b we show *m*0_*min*_ as a function of *α* for *f* = 0.01 and *θ* = 0.4,0.6 and 0.8. For a given value of *θ*, *m*0_*min*_ is only weakly modulated by the storage load, with *m*0_*min*_ ≃ *θ*. This can be seen from equations (7), which give, for small *f*,

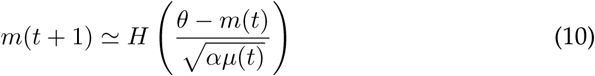

When *αμ*(*t*) ≪ 1 we have

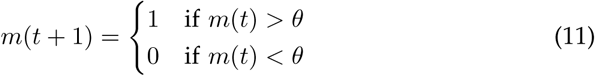

## 4 Memory properties of a single path

After isolating a single module from a TRN, we isolate a single feed-forward path composed of *L* modules, from the root module up to one of the leaves of the tree. Stimuli propagate along the feed-forward structure from the root module. We first describe the propagation of stimuli orthogonal to the stored patterns. This allows us to measure the buffering capacity, which quantifies the ability of the neural activity of the path to represent a sequence of previously observed stimuli (Fig. 4a). We then consider a stimulus that is stored in one of the module, and describe under which conditions it can trigger a state of persistent activity in this module, as illustrated in Fig. 5a. This allows us to quantify the storage capacity of a path of length *L*.

### 4.1 Dynamical equations for propagation of activity patterns

Here modules form a single feed-forward path and are indexed with *l* = 1…*L*. A stream of sensory stimuli imposes the state of the root module ***σ***^0^ at all times with

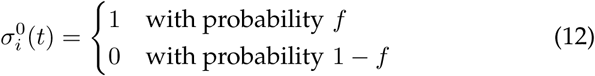

An input represented in the root module at time *t*−*l* reaches module *l* at time *t*. The dynamics of the path is characterized by {*μ*^*l*^(*t*)}_*l*=1…*L*_ the mean activities in the different modules, and {*n*^*l*^(*t*)}_*l*=1…*L*_ the overlaps between ***σ***^0^(*t*−*l*) and ***σ***^*l*^(*t*)

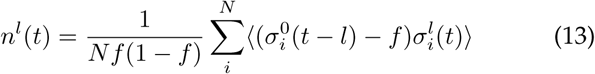

The time evolution of these macroscopic variables is governed by (see Supplementary materials for derivation)

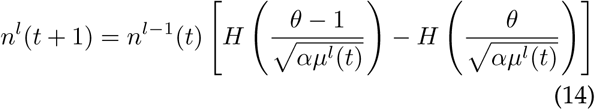

together with

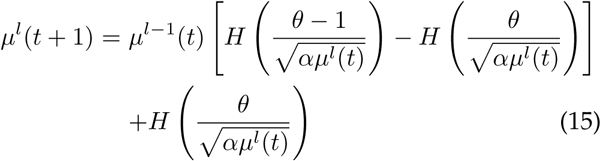

### 4.2 Mean activity in steady state

To understand how stimuli propagate throughout the path, it is informative to first examine the profile of mean activity the path settles into under the influence of the stream of stimuli. At the macroscopic level this profile is given by the steady states of (14)(15).

At the microscopic level a neuron i in module *l* is active if

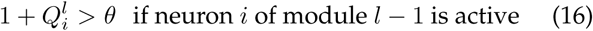

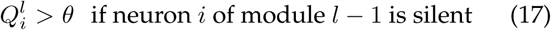

where 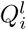 corresponds to the recurrent noise coming from the storage of patterns in module *l*.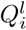 is drawn from a normal distribution 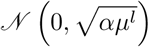. Depending on the value of *θ* and the standard deviation 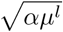, two behaviors are possible, either the average activity in each module, *μ*^*l*^, increases along the path (regime I), either it decreases (regime II).

In the first regime the increase is characterized by a critical depth *L*_*c*_ from which activity explodes in the network: *μ*^*l*<*L*_*c*_^ = *O*(*f*) and *μ*^*l*>*L*_*c*_^ ≫ *f*. The *α* − *θ* region of parameters in which this behavior is observed is shown in orange in Fig. 3a. The frontiers of this region can be derived analytically for small coding levels *f* ≪ 1 (see Appendix 7.1). It corresponds to low activation thresholds 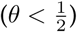 or high activation threshold paired with high recurrent noise (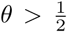 and *α* > (2*θ*^−1^ − *θ*^−2^)*α*_*c*_). An example of profile of average activity *μ*^*l*^,*l* = 1…*L* is shown in the top panel of Fig. 3b. Values of *L*_*c*_ can also be derived analytically (see Appendix 7.1) and its dependence on *α* and *θ* are shown in Fig. 3d. As the storage load *α* is increased the recurrent noise becomes more effective at activating neurons and the explosion of activity occurs at modules closer to the root module (top panel). As the activation threshold *θ* is increased, it becomes more difficult for the recurrent noise to activate neurons and *L*_*c*_ increases (bottom panel).

**Figure 3.**
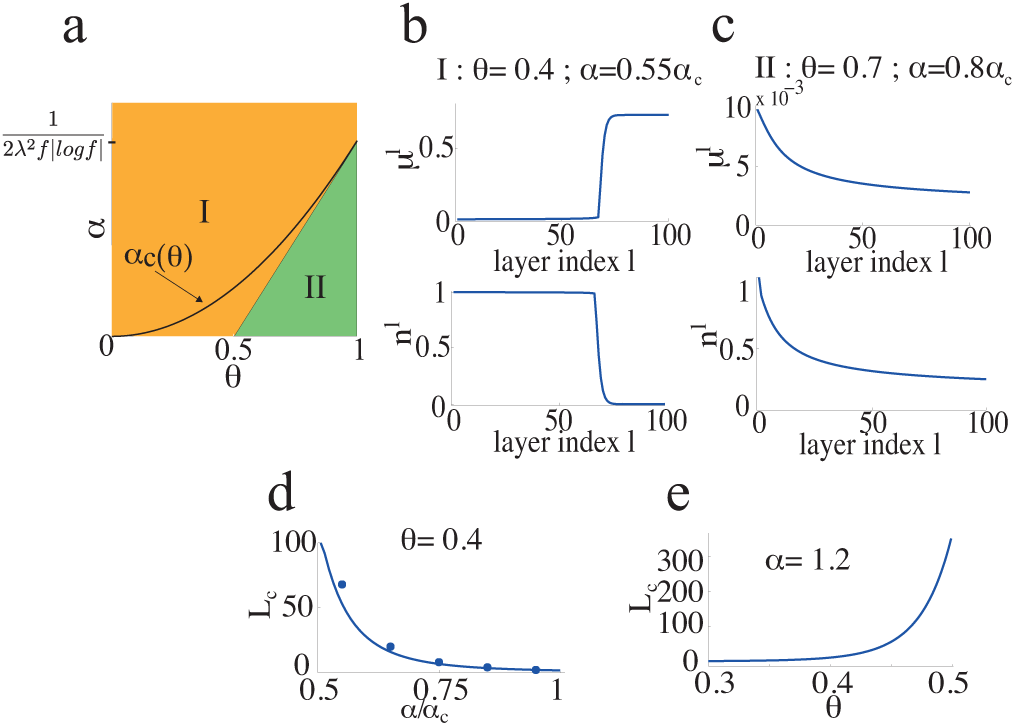
Propagation of stimuli throughout the path. **a** Mean activity in a path. Regions in the θ − α plane where the different regimes are found for f ≪ 1. In regime I, the orange region above the oblique line, μ^*l*^ is increasing with l. In regime II, green region below the oblique line, μ^*l*^ is decreasing with l. The parabolic line represents the storage capacity α_*c*_. **b** Top: average activities in steady state for a path of L = 100 modules. Bottom: overlap n^*l*^ between a pattern sent at time t and the state of module l at time t + l. **c** Same as b for regime II. **d** L_*c*_ as a function of the normalized storage load 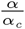 for θ = 0.4. e L_c_ as a function of θ for α = 1.2.

**Figure 4.**
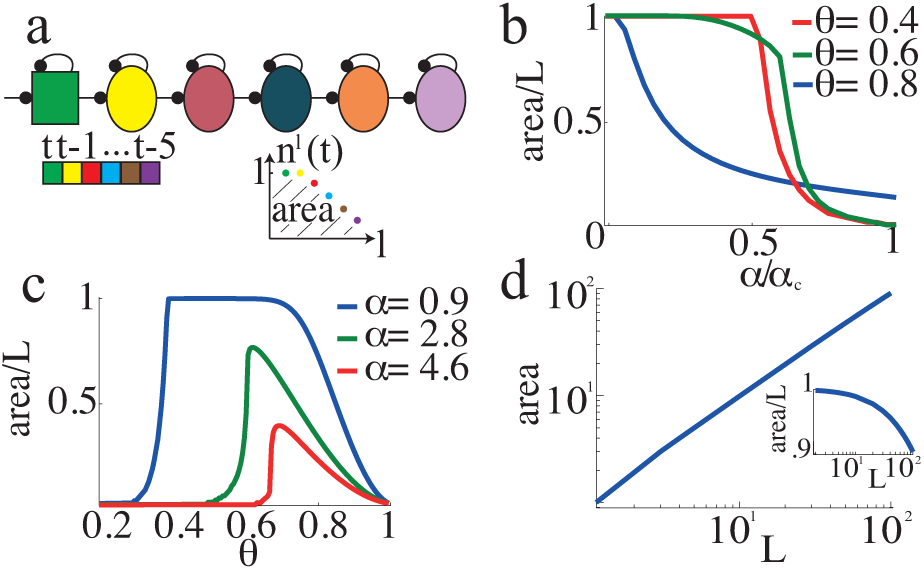
Buffering of arbitrary input sequences. **a** Illustration of a path buffering the sequence of stimuli: ‘magenta,brown,blue,red,yellow,green’, with the ‘green’ stimulus presented at time t in the root module. The buffering capacity of a path of length L is measured by the area under the curve n^l^, measuring the overlap between the activities in individual modules and patterns of activity previously presented in the root module. b Area as a function of the amount of recurrent connections α/α_*c*_(*θ*) for L = 100. **c** Normalized buffering capacity as a function of θ, L = 100. **d** Buffering capacity as a function of path length for θ = 0.6 and α = 2. Insets shows how the normalized buffering capacity slightly decreases with L.

**Figure 5.**
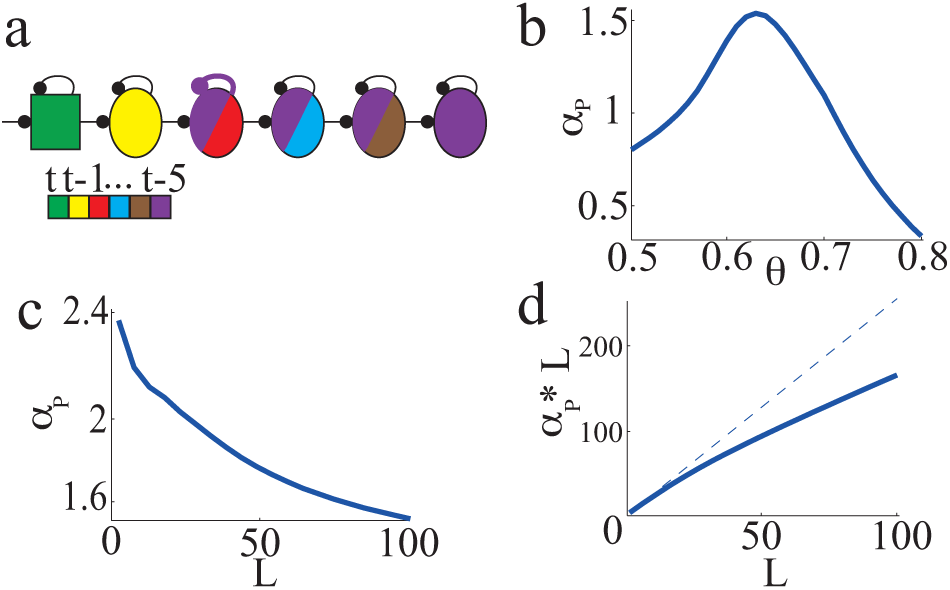
Storage capacity of a path. **a** Illustration of retrieval in a path, a sequence of stimuli ‘magenta,brown,blue,red,yellow,green’ is presented by the root module. The recurrent connectivity of the second module is designed to maintain the magenta stimulus in short-term memory. Once retrieval has been elicited in this module, the activity in this module and subsequent ones represent the magenta stimulus together with the ongoing sequence. **b** Optimal single module storage load as a function of θ for a path of length L = 100. **c** Optimal single module storage load as a function of path’s length. d Sub-linear increase of storage capacity as a function of path’s length. Storage capacity is optimized over θ for each value of L here.

In the complementary region of parameter space (green in Fig. 3a), the mean activity *μ*^*l*^ decreases with *l*. In this regime the recurrent noise mainly affects the mean activity with a negative input. An example of *μ*^*l*^,*l* = 1…*L* in this case is shown in the top panel of Fig. 3c.

### 4.3 Short-term memory via transient activity in a path

In this section we quantify the ability of a path to maintain a memory of a sequence of stimuli in its transient neural activity, i.e. its buffering capacity. When there are no recurrent connections (*α* = 0), inputs propagate through the path without being corrupted, with the input sent at time *t* by the root module being perfectly represented in module *l* at time *t*+*l* (i.e. *n*^*l*^ (*t*) = 1 for all *l*, *t*). The transient state of the path thus buffers memory sequences that last *L* time steps, where *L* is the total number of modules in the path. In the presence of recurrent connections feed-forward inputs carrying the stimuli are in competition with recurrent inputs acting as random noise (because stimuli are uncorrelated with 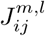). Stimuli are thus progressively degraded as they travel along the path, according to (14). This is illustrated in the bottom panels of Fig. 3b-c, where we show *n*^*l*^ in the different regimes of mean activity described above. In regimes I, patterns are slightly degraded by their travel through each module, until they reach module *l* ≥ *L*_*c*_ where *n*^*l*>*L*_*c*_^ drops to a value close to zero. Before reaching this critical module, the overlap decreases moderately since degradation mainly occurs by activating silent neurons, which does not dramatically affect the overlaps for the small coding levels considered here. In regime II, the overlap decreases with *l*, smoothly but relatively fast, since silencing neurons that should be active has a strong impact on the value of the overlaps. In order to quantify the ability of a path to buffer inputs, we define the buffering capacity as the area under the *n*^*l*^ versus *l* curve (White et al., 2004). In Fig. 4b we show how this area decreases with the amount of recurrent connections for different values of *θ*, and *L* = 100. For *θ* = 0.4, the profile of mean activity corresponds to regime I and the area under the *n*^*l*^ curve is close to *L* (*L* inputs are perfectly represented in the path of length *L*) until *L*_*c*_(*α*) becomes smaller than *L* and the area decreases sharply with the amount of recurrent connections. For *θ* = 0.6 and 0.8 the observed decrease occurs in regime II. The specific effect of *θ* on the buffering capacity is shown in Fig. 4c. The optimal value of *θ* varies from 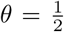 at *α* ≃ 0 to *θ* ≃ 0.65 at *α* = 4.6, the storage capacity of a single module. Finally in Fig. 4d, we show how the buffering capacity scales with *L* for fixed parameters *θ* = 0.6 and *α* = 2.

### 4.4 Short-term memory via reverberating activity in a path

In this section the root module conveys a stream of random stimuli, except at *t* = 0 for which the root module is set in the state ***σ***^0^(*t* = 0) = **ξ**^*μ*_0_,*l*_0_^ corresponding to a pattern stored in the synaptic matrix of module *l*_0_. This scenario is illustrated in Fig. 5a. To describe the propagation and retrieval of this pattern we introduce *m*^*l*^ (*t*), the overlap between ***σ***^1^(*t*), the state of module *l* at time *t*, and ***ξ***^*μ*_0_,*l*_0_^

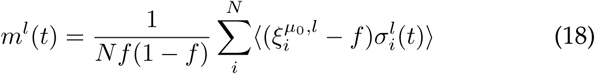

In module *l* < *l*_0_, pattern ***ξ***^*μ*_0_,*l*_0_^ has the same status than a random stimulus, and the order parameter *m*^*l*<*l*_0_^(*t*) is governed by equation (14). In module *l* = *l*_0_, recurrent connections tend to sustain the pattern ***ξ***^*μ*_0_,*l*_0_^, and the overlap obeys equation (28) which can be found in the Appendix 7.2. Once the pattern is retrieved in module *l*_0_, it propagates further in deeper modules *l* > *l*_0_.

Retrieval in module *l*_0_ differs from the single module case in two aspects: first, the retrieval state has to persist despite the presence of feed-forward noise caused by random patterns of activity that enter the path at *t* ≠ 0. And second, at *t* = *l*_0_, module *l*_0_ is cued with a noisy version of ***ξ***^*μ*_0_,*l*_0_^. Below we study these two effects separately and then show how they impact the storage capacity of a path.

#### 4.4.1 Effect of feed-forward inputs on the existence of a retrieval state

The ongoing stream of stimuli in the path tends to destabilize a retrieved pattern by activating neurons that should not be active. This increase of activity (*μ*^*l*^ ≃ 2*f* in modules close to the root module) mediates an increase in recurrent noise since the recurrent connections also reflect the storage of patterns ***ξ***^*μ*,*m*_0_^ ≠ ***ξ***^*μ*_0_,*m*_0_^. In order to accommodate this higher recurrent noise, modules embedded in a path need to have a storage load smaller than *α*_c_. The specific condition is given by (see Appendix 7.3 for derivation):

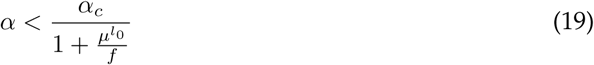

where *μ*^*l*_0_^ is the mean activity in module *l*_0_ induced by the random stream of inputs, before retrieval, which is obtained from the steady-state solution of equation (15), and *α*_*c*_ is the storage capacity of an isolated module. For modules close to the root module for which *μ*^*l*_0_^ ≃ *f*, equation (19) leads to a maximal storage load 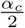 under the presence of feed-forward noise. In deeper modules, the value of the maximal storage load for which the retrieval state is stable depends on the profile of mean activity *μ*^*l*^, *l* = 1…*L*. In regime I (Fig. 3a), for modules close to the critical module *l ≲ L*_*c*_, where the mean activity is larger than *f*, the maximal storage load will be even more reduced. In regime II, the mean activity decreases with *l* and the maximal storage load compatible with the feed-forward drive can in principle be higher than for the initial module. Note however that we do not take advantage of this effect in the present work, since for simplicity we only consider equal storage load in each module.

#### 4.4.2 Retrieval from corrupted signal

The signal eliciting retrieval in module *l*_0_ is a corrupted version of pattern ***ξ***^*μ*_0_,*l*_0_^, i.e. *m*^*l*_0_^(*t* = *l*_0_) = *m*_0_ < 1. The success of retrieval thus depends on whether this corrupted version is within the basin of attraction of ***ξ***^*μ*_0_,*l*_0_^. Similarly to the single module case, assuming *μ*^*l*_0_^ = *O*(*f*) and *f* ≪ 1, the condition for retrieval is:

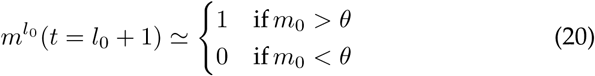

#### 4.4.3 Storage capacity of a path

Because of the two phenomenons described above, modules of the TRN should have a storage load lower than the storage capacity of an isolated module (*α* < *α*_*c*_) in order for a path to serve as an auto-associative memory. We have iterated the dynamical equations (28) (see Appendix 7.2) in order to measure the storage capacity of paths of length *L*

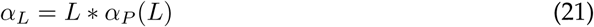

Where *α*_*L*_ quantifies the total number of patterns that can be retrieved by cueing from the root module. In the definition of *α*_*L*_ we also impose that retrieved memories can be read-out in the last module of the path, which we define as 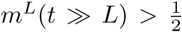 (see Discussion). *α*_*L*_ crucially depends on the parameter *θ* because i) it controls the size of the basins of attractions, cf (20); ii) the single module storage load *α*_*c*_ depends on *θ*; and iii) the corruption of patterns during feed-forward propagation strongly depends on *θ* (e.g. Fig. 4c). In Fig. 5b we show the storage capacity *α*_*L*=100_ as a function of *θ*, a maximum of *α*_*L*_ ≃ 0.3 _* *α*_*c*_ * *L*_ is reached for *θ* ≃ 0.6. In other words, to accommodate all the constraints related to having an attractor network embedded in a feed-forward path of length *L* = 100, the storage load of each module 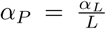 has to be reduced by ≃ 1/3 compared to the case of an isolated module. This ratio decreases further as the length of the path increases (Fig. 5c), going from a ratio of 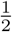 for *L* = 1 (maximal ratio that allows to sustain a pattern with a mean activity 2*f* in the presence of feed-forward inputs aacording to (19)) to ≃ 0.23 for *L* = 50, 000 (not shown). Figure 5d shows how the capacity of a path *α_L_* increases sub-linearly with *L*.

## 5 Memory properties of the full TRN

As seen in the previous section, the memory properties of a path crucially depends on its length *L*. The buffering capacity scales with the length of the path, if working with appropriate parameters, e.g. avoiding explosion of mean activity with storage load small enough to have *L*_*c*_ > *L*. And the maximal storage load per module *α*_*P*_(*L*) decreases with *L* due to increasing noise accumulation with paths lengths. We now use these results to describe the memory properties of full TRNs of *M* modules. TRN are composed of feed-forward paths of length *L*, where *L* is related to *M* through the divergence ratio *d*, measuring the ratio between the numbers of modules in two successive levels of the TRN (see Fig. 1):

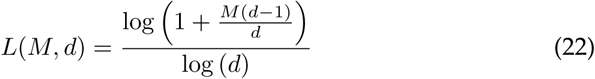

### 5.1 Storage capacity of TRN

The TRN being composed of feed-forward paths of length *L*, its storage capacity is given by

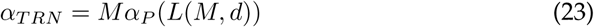

as defined above *α*_*P*_ (*L*) is the maximal storage load of a single module that ensures proper working of a path of length *L*. As shown in Fig. 5b, *α*_*P*_(*L*) is a decreasing function of *L*, and as such the capacity of the TRN is maximized for a fully parallel architecture *L* = 1 (*d* = *M*) and minimized for a fully serial architecture *L* = *M* (*d* = 1). This is shown in Fig. 6 where we show *α*_*TRN*_ as a function of d for a TRN of *M* = 50, 000 modules, which would for instance correspond to a sensory-hierarchy of 500 * 10^6^ neurons with columns composed of 10^4^ neurons. From the fully serial to the fully parallel organization, the total capacity *α_TRN_* is increased by a factor ≃ 3.

### 5.2 Buffering inputs in the TRN

The ability of the TRN to buffer inputs is strongly dependent on the length of its paths, as shown in Fig. 4d. In principle the buffering capacity of a TRN of depth *L* is larger than the buffering capacity of a path of length *L*, since the same patterns are represented in multiple paths and this redundancy could be exploited by a readout. On the other hand the buffering capacity is upper bounded by the length of the paths *L*. This lower and upper bounds correspond to the cyan and blue curves in Fig. 6b, which decreases sharply with *d.*

**Figure 6.**
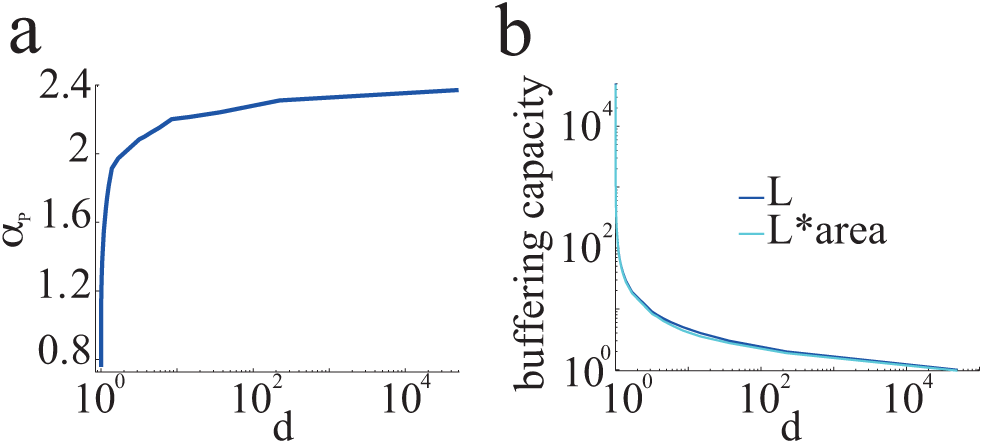
Memory properties of full TRNs of M = 50, 000 modules. On the x-axis d = 10° corresponds to a fully serial architecture as shown in Fig. 1a, while d = M corresponds to a fully parallel architecture as shown in Fig. 1b. **a** Storage capacity as a function of the expansion ratio. **b** Lower and upper bounds (cyan and blue curves) on the buffering capacity as a function of the expansion ratio, for θ = 0.4, α = 0.85.

## 6 Conclusion

In this paper, we have studied the short-term memory properties of trees of recurrent networks (TRN). For this neural architecture the feed-forward tree-like backbone of modules supports short-term memory via transient dynamics (Ganguli et al., 2008; Jaeger and Haas, 2004; Lim and Goldman, 2012; Maass et al., 2002; White et al., 2004), and the recurrent connectivity inside each module supports short-term memory via reverberating activity (Amit, 1989; Amit and Brunel, 1997; Hopfield, 1982; Tsodyks and Feigel*’*man, 1988). The ability of TRN to memorize sequences of stimuli via transient dynamics has been quantified using the buffering capacity (White et al., 2004), a measure of how many stimuli, successively presented at the first module, are well represented in the pattern of activity of the TRN at a given time. Networks with feed-forward structures have been shown to be optimal to implement such memory mechanisms, since they avoid interference between patterns presented at different times (Ganguli et al., 2008; Lim and Goldman, 2012). In TRN, stimuli propagate in a feed-forward manner and reach modules of increasing depth as time is incremented. The buffering capacity of a TRN is thus circumscribed by its depth, and crucially depends on a faithful propagation of stimuli along the network. We have shown that such a faithful propagation is mainly controlled by two factors. The amount of recurrent connections in each module, controlling the strength of the recurrent inputs elicited by a given level of activity in each module; those inputs act as noise by competing with the feed-forward inputs. The activation threshold of neurons, which sets the levels of activity in the modules, as well as tunes signal corruption, by controlling whether recurrent noise tends to activate neurons that should be silent or active according to the incoming feed-forward input.

The ability of TRN to hold a stimulus in memory via reverberating activity has been quantified with the storage capacity, i.e. the number of patterns of activity that modules can maintain in persistent activity, see e.g. (Amit, 1989). Compared to standard studies of an isolated attractor network, the capacity analysis of TRN requires to take into account two additional constraints. First, a state of persistent activity in a module needs to be robust to feed-forward inputs that act as a source of noise. The consequence of this noise is an increased mean activity in the module, leading to more recurrent noise. This requires to decrease the amount of stored patterns per modules compared to the case of an isolated network. Second, the network is cued with a corrupted signal, due to noisy propagation of signals along a TRN. This requires to be able to compute the extent to which patterns of activity are corrupted when traveling through a TRN, as well as to compute the size of the basins of attractions associated to each stored memory pattern. Again the activation threshold plays here a crucial role, since it controls signal propagation, the size of the basins of attractions, as well as the single module storage capacity. Because a pattern of activity is all the more corrupted as it has travelled through many modules, TRN with low depth are better suited to reach high storage capacities.

TRNs are able to support both kinds of short-term memory mechanisms, with the non trivial observation that similar values of a key parameter, the activation threshold, are optimal in both settings. Although both kinds of short-term memory are optimized by similar values of the activation threshold, the buffering and storage capacity of a TRN depend strongly on its depth, with a purely serial TRN architecture being optimal for short term memory via transient dynamics and a purely parallel architecture being optimal for short-term memory via reverberating activity.

We have introduced TRN in order to investigate memory properties of sensory cortices, which can be described as made of a few neurons driven by sensory stimuli via thalamic inputs, and a vast majority of interneurons mainly receiving cortical inputs (Braitenberg and Schütz, 1991). Cortico-cortical connectivity has been shown to be spatially clustered (Bosking et al., 1997; DeFelipe et al., 1986; Gilbert and Wiesel, 1989; Pucak et al., 1996), leading to the notion of cortical column. These columns are modeled by the modules of our network, which we have taken to be attractor neural networks, a class of networks whose connectivity profile is consistent with local connectivity in sensory cortices (Brunel, 2016), and that has been shown to efficiently encode sensory stimuli (Tkačik et al., 2010). The extent to which cortical columns are organized in series or in parallel with respect to the thalamic inputs is not known, although, in the visual hierarchy, available data makes it clear that there is some form of serial feed-forward organization. Anatomy has shown that as the depth in the hierarchy is increased, networks are less and less under the influence of the thalamus, with ≃ 0.2% of inputs to V1 neurons having a thalamic origin, versus ≃ 0.1% for V2 and ≃ 0.05% for V4 neurons (Markov et al., 2010). It has also been shown that activity in V2 is strongly dependent on activity in V1, as suggested by lesions of V1 suppressing activity in V2 (Girard and Bullier, 1989). Moreover, along the visual hierarchy, stimuli are processed with increasing delays, with 40 ms and 80 ms delays in V1 and infero-temporal cortex respectively, suggesting a form of serial processing (Thorpe and Fabre-Thorpe, 2001). However, whether such serial organization supports short-term memory via transient dynamics is not obvious since this difference in processing delays between V1 and infero-temporal cortex is much smaller than the behavioral time scales associated with short-term memory. Nevertheless it can be argued that more advanced analysis could reveal short-term memory via transient dynamics at the scale of a brain area (Klampfl et al., 2012; Nikolić et al., 2009).

Neural activity in a TRN forms sensory representations of ongoing stimuli that can be held in short-term memory. A question we have not discussed so far is the one of the reading of such representations by a read-out system. For short-term memory with reverberating activity, once a stimulus has elicited a pattern of persistent activity in one of the module, this pattern propagates down to sub-sequent modules (see Fig. 5a). A natural read-out for such a representation would be the last module of each path, which would correspond to associative areas in a cortical hierarchy. In Supplementary materials we show how reading-out through the last module of a path is done at the expense of reducing the storage load to avoid dramatic signal degradation from the module maintaining persistent activity to the end of the paths. Regarding short-term memory with transient dynamics, which buffers sequences of stimuli encoded on the whole depth of a TRN, a read-out system has to be connected to modules of all depths in order for reading such sensory representations (White et al., 2004). In a cortical setting such a read-out system could also be implemented by areas on the top of sensory hierarchies, with feed-back connections, not modeled in a TRN, sampling from areas lower in the hierarchy.

Anatomical studies reveal a cortical architecture that is much richer than the simple architecture of TRN, which include neither feed-back nor shortcuts between modules. Other simplifications are the one-to-one projections from one layer to the next, and the fact that patterns stored in different layers have no semantic relationships. While these simplifications allowed us to quantify the memory properties of such networks in depth, it remains to be investigated how these properties will be affected in more realistic networks. Nevertheless it is remarkable that it has remained possible to track analytically the dynamics of such cellular-resolution large-scale networks, and evaluate the computational performance of various connectivity schemes, such as serial or parallel. Such an approach, linking structural and functional properties of this type of network models, together with the advent of large-scale single-cell resolution connectomes, will allow to shed new lights on the computational roles of cortical networks.

## 7 Appendix

### 7.1 Steady-state mean activity profiles for *f* ≪ 1

We consider a module with mean activity *μ* = *O*(*f*) receiving feed-forward inputs from a module whose mean activity is 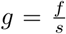. From equations (15), the fixed point equations relating *μ* and *g* is

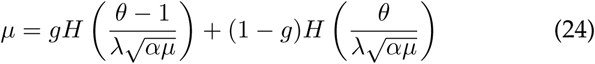

Using the estimate 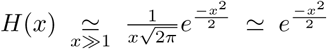, and 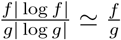 the the fixed point equation can be rewritten

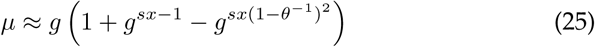

with 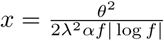. Comparing the two last terms of equation (25) allows to understand whether *μ* increases or decreases compared to *g* and thus to describe the two regimes of mean activity profiles of section 4.2. Moreover, for the regime of increasing activity along the path, the transition from *μ* = *O*(*f*) to *μ* = *O*(1) arises for *s* = 1/*x*, i.e. *g* = *xf*. By differentiating (24) as shown in Supplementary materials, this allows to give expressions for the depth *L*_*c*_ at which the transition occurs. For 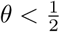

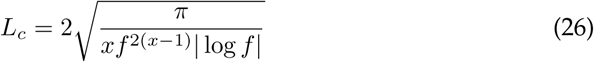

and for 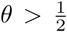 and *α* > (2*θ*^−1^ −*θ*^−2^)*α*_*c*_ the critical depth scale as

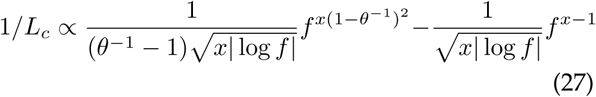

### 7.2 Dynamical equations for memory retrieval in a path

In order to describe the retrieval of pattern *ξ*^*l*_0_,1^ in a module *l*_0_ receiving feed-forward inputs from module *l*_0_ − 1 (see section 4.4), we have used the following dynamical equations, whose derivation is detailed in Supplementary Materials. *m*^*l*_0_^ (resp. *m*^*l*_0−1_^) is the overlap between the activity in module *l*_0_ (resp. *l*_0_ − 1) and *ξ*^*l*_0_,1^.

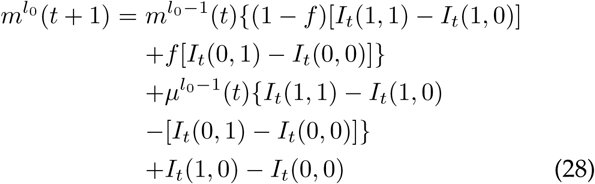

and

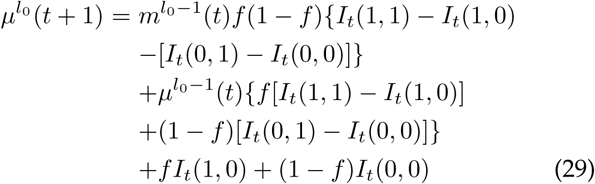

with

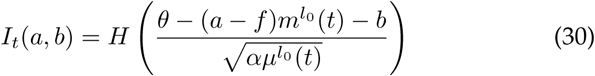

### 7.3 Impact of feed-forward noise on persistent activity

The random sequence of inputs is a form of noise that reduces the capacity for retrieval states. To evaluate the existence of a retrieval state *ξ*^*μ*_0_,*m*_0_^ under these conditions, we examine the stability of the module *l* assuming that it receives random feed-forward inputs from the module *l* − 1 with a steady state mean activity 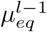. We re-write the equations for retrieval in module *l* in terms of the order parameters 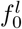 and 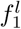, which measure the fraction of background neurons (neurons *i* such that 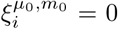) that are active and the fraction of foreground (neurons *i* such that 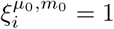) neurons that are active. These order parameters are related to *m*^*l*^ and *μ*^*l*^ by 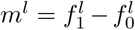 and 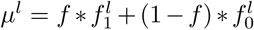.

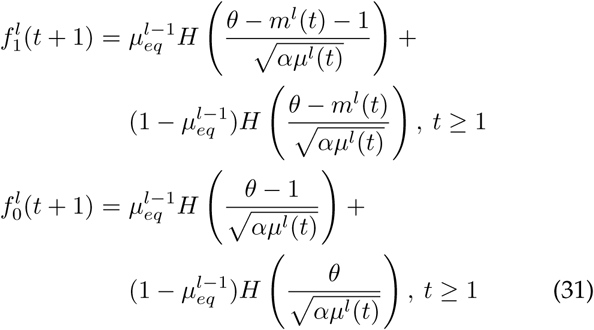

If we assume that a pattern has been retrieved, i.e. *m*^*l*^(*t* = 1) ≃1 and 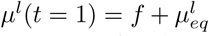. This pattern remains stable if 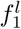 remains of order 1 and 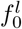 remains of order *f*, which comes down to have, for 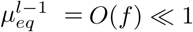,

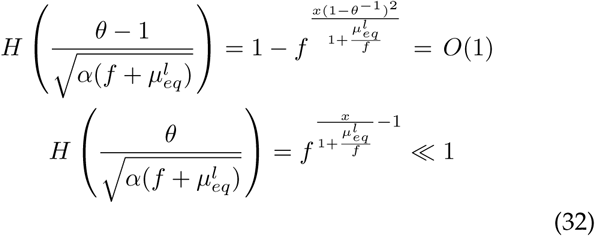

For *f* ≪ 1 this is satisfied if 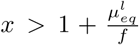, hence the condition (19) on *α*.

## Acknowledgements

I would like to thank Nicolas Brunel and Haim Sompolinsky for useful discussions throughout the course this work.

