## Supplementary materials for "Short term memory properties of sensory neural architectures"

#### 1 A more general model

In the more general model of this document, a parameter $\lambda$ scales the ratio of recurrent inputs and feed-forward inputs.

$$h_i^m(t) = \sigma_i^{a(m)}(t) + \lambda \sum_{j=1}^{N} J_{ij}^m \sigma_j^m(t) \tag{1}$$

where module $a(m)$ is the ancestor of module $m$.

##### 1.1 Dynamics at T>0

In the main text the network evolution is deterministic, but the study can be extended to stochastic dynamics by considering the following Glauber dynamics, parametrized by a temperature $T$

$$\sigma_i^m(t+1) = \begin{cases} 1 & \text{with probability}(1+\exp[-\frac{h_i^m(t)-\theta}{T}])^{-1} \\ 0 & \text{with probability}(1+\exp[\frac{h_i^m(t)-\theta}{T}])^{-1} \end{cases} \tag{2}$$

##### 1.2 Diluted connectivity

Patterns are stored by a randomly diluted recurrent connectivity matrix, using a covariance learning rule [Sejnowski, 1977, Tsodyks and Feigel'man, 1988]

$$J_{ij}^l = \frac{C_{ij}^l}{Cf(1-f)} \sum_{\mu=1}^{P} (\xi_i^{\mu,l} - f)(\xi_j^{\mu,l} - f) \tag{3}$$

where $C_{ij}^l$ is a random asymmetric adjacency matrix, such that:

$$Pr(C_{ij}^l = 1) = \frac{C}{N} \text{ and } Pr(C_{ij}^l = 0) = (1 - \frac{C}{N}) \tag{4}$$

and C is the connection probability. We introduce the memory load $\alpha = P/C$ which quantifies the number of memories stored in each layer.

### 2 Single module

#### 2.1 Derivation of the dynamical equation

In what follows, we use the reasoning of Derrida [Derrida et al., 1987], applied to our model. We realize this has been done previously in [Evans, 1989]. In the limit of extreme dilution ($C \ll \ln N$) it allows to rigorously derive equations for the dynamic of the order parameters. These equations have the same fixed points than those derived by Tsodyks [Tsodyks and Feigel'man, 1988] in the limit $C \ll N$, or $C = N$ and small $f$. Simulations suggest that the dynamical equations can be extrapolated to non-diluted networks.
We will work in the limit $N \to \infty$. In order to study the retrieval of a single memory we characterize the state of the network using two macroscopic variables:

$$m(t) = \frac{1}{Nf(1-f)} \sum_i^N \langle (\xi_i^1 - f)\sigma_i(t) \rangle \tag{5}$$

and

$$\mu(t) = \frac{1}{N} \sum_i^N \langle \sigma_i(t) \rangle \tag{6}$$

where $m(t)$ is the overlap between the state of the network and the memory $\vec{\xi}^1$ and $\mu(t)$ is the fraction of active neurons in the network at time $t$. Averages are taken over different microscopic initial states and over network's dynamics.

If we consider a site $i$ that is connected to a tree of K presynaptic neurons, and if we are interested in the overlap with the first pattern $\vec{\xi}^1$, the field $h_i(t)$ can be written

$$h_i(t) = \frac{1}{Cf(1-f)}(\xi_i^1 - f)\sum_{r=1}^{K}(\xi_{j_r}^1 - f)\sigma_{j_r}(t) + \frac{1}{Cf(1-f)}\sum_{\mu=2}^{p}(\xi_i^\mu - f)\sum_{r=1}^{K}(\xi_{j_r}^\mu - f)\sigma_{j_r}(t) \tag{7}$$

At fixed C, such a tree of presynaptic neurons appears with probability $\frac{C^K e^{-K}}{K!}$, so that the average length and the most probable length for a tree is C. In what follows we will consider the limit $C \to \infty$, and replace K by C, i.e. we average $h_i$ over the different possible shapes of the tree of presynaptic neurons. The first term of $h_i$ can be written:

$$\begin{aligned} s_i(t) &= \frac{1}{Cf(1-f)}(\xi_i^1 - f)\sum_{r=1}^{C}(\xi_{j_r}^1 - f)\sigma_{j_r}(t) \\ &= (\xi_i^1 - f)(m(t) + O(\frac{1}{C})) \end{aligned} \tag{8}$$

Let us now consider the second term in $h_i(t)$. If $C \ll \log N$, then the variables $\sigma_{j1}(t), ..., \sigma_{jC}(t)$ are uncorrelated and the second term of $h_i(t)$ can be treated, for large P, as a gaussian noise $z_i(t)$ of mean 0 and variance $\sigma_z = \frac{p}{C}\mu(t)$. In this limit the noise in the first term whose variance is of order $O(\frac{1}{C})$ is negligible compare to this noise term whose variance is of order

$\alpha\mu(t)$ (with $\alpha = \frac{p}{C}$).
So we can rewrite:

$$h_i(t) = (\xi_i^1 - f)m(t) + z_i(t) \quad (9)$$

We can now turn to the computation of $m(t+1)$:

$$\begin{aligned} m(t+1) &= \frac{1}{Nf(1-f)} \sum_i \langle (\xi_i^1 - f)\sigma_i(t+1) \rangle & (10) \\ &= f_1(t+1) - f_0(t+1) & (11) \end{aligned}$$

where $f_1(t+1) = \langle \sigma_i(t+1) \rangle_{\xi_i^1=1}$ (*resp.*$f_0(t+1) = \langle \sigma_i(t+1) \rangle_{\xi_i^1=0}$) is the average of $\sigma_i(t+1)$ for sites i such that $\xi_i^1 = 1$ (*resp.* $\xi_i^1 = 0$). So one can express $m(t+1)$ as a function of $m(t)$ and $\mu(t)$:

$$\begin{aligned} m(t+1) &= \int_{-\infty}^{+\infty} \frac{dz}{\sqrt{2\pi}} e^{-\frac{z^2}{2}} \left[ \frac{1}{1 + \exp(-\frac{1}{T}[(1-f)m(t) + \sqrt{\alpha\mu(t)}z - \theta])} \right. \\ & \left. - \frac{1}{1 + \exp(-\frac{1}{T}[-fm(t) + \sqrt{\alpha\mu(t)}z - \theta])} \right] & (12) \end{aligned}$$

separating sites for which $\xi_i^1 = 1$ and $\xi_i^1 = 0$, one can show that $\mu(t+1)$ is given by:

$$\begin{aligned} \mu(t+1) &= \int_{-\infty}^{+\infty} \frac{dz}{\sqrt{2\pi}} e^{-\frac{z^2}{2}} \left[ \frac{f}{1 + \exp(-\frac{1}{T}[(1-f)m(t) + \sqrt{\alpha\mu(t)}z - \theta])} \right. \\ &+ \left. \frac{1-f}{1 + \exp(-\frac{1}{T}[-fm(t) + \sqrt{\alpha\mu(t)}z - \theta])} \right] & (13) \end{aligned}$$

#### 2.2 Capacity analysis at $T = 0$

Here we recapitulate results of previous analyses [Tsodyks and Feigel'man, 1988, Sompolinsky and White, 2003] that lead to the characterization of the strong retrieval state. We first study the existence of a fixed point with high overlap with one of the memories, say $\vec{\xi}^1$. For small enough $\alpha$, and $0 < \theta < 1$, we can write,

$$\begin{aligned} 1 - f_1 &= H\left(\frac{-\theta + m}{\sqrt{\alpha\mu}}\right) \approx \exp\left(-\frac{(1-\theta)^2}{2\alpha\mu}\right) \ll 1 & (14) \\ f_0 &= H\left(\frac{\theta}{\sqrt{\alpha\mu}}\right) \approx \exp\left(-\frac{\theta^2}{2\alpha\mu}\right) \ll 1 & (15) \end{aligned}$$

These equations have to be solved self-consistently using,

$$\mu \approx f(f_1 + f_0/f) = f + f(f_0/f - (1 - f_1)) \quad (16)$$

For a retrieval state to exist, not only $1 - f_1$ and $f_0$ must be small, but also $f_0/f$ must be small, so that the number of active neurons with $\xi_i^1 = 0$ is much smaller than that of the correctly active neurons. When this is the case (i.e., for $\alpha$ below capacity)

$$\begin{aligned} m &\approx 1 \\ \mu &\approx f \end{aligned}$$

The capacity is reached when $f_0$ increases sharply so that $f_0$ becomes of order $f$ causing a sharp increase in mean activity level $\mu$, which also implies a large increase in the noise of the field, taking the system away from the memory state. To obtain a quantitative analysis of the capacity, we introduce the following scaling

$$\alpha = \frac{\theta^2}{2xf|\log f|} \tag{17}$$

and consider the behavior of the system as $x$ increases. Substituting in equations (14)-(17), we obtain,

$$1 - f_1 \approx f^{x(1-\theta^{-1})^2} \tag{18}$$

$$f_0/f = f^{(x-1)} \tag{19}$$

These errors are small as long as $x > 1$, so that the maximum capacity is given by $x = 1$, i.e., $\alpha < \frac{\theta^2}{2f|\log f|}$. This result holds in the limit of small $f$. For moderate values of $f$ the capacity increases initially quadratically with $\theta$ but beyond some $\theta_C$ it decreases until it vanishes as $\theta \to 1$. We can estimate $\theta_C$ by demanding that the

$$-\log(1 - f_1) \approx \frac{(1-\theta)^2}{2\alpha\mu} \tag{20}$$

(see (14)) be larger than 1. Substituting (17) and reintroducing $\lambda$ (and $0 < \theta < \lambda$ ) yields $x(1 - \lambda\theta^{-1})^2 > 1/|\log f|$. Thus, we can write

$$x > \max[1, \frac{1}{(1-\lambda\theta^{-1})^2|\log f|}] \tag{21}$$

and the equation for the capacity

$$\alpha_C = \min\left[\frac{\theta^2}{2\lambda^2 f|\log f|}, \frac{1}{2f}(1-\lambda^{-1}\theta)^2\right] \tag{22}$$

and $\theta_0$ where the two branches meet is given by

$$\theta_0 = \frac{\lambda}{1 + \frac{1}{\sqrt{|\log f|}}} \tag{23}$$

**Activity levels below the transition:**
It is interesting to ask what is the main source of deviation of $\mu/f$ from 1 below capacity: decrease in $\mu$ by $1 - f_1$ or increase due to $f_0/f$ ? As shown above, close to the capacity, the dominant term is an increase in $\mu$ due to $f_0/f$. However, this is not necessarily the case below the transition. From equations (18),(19), the condition for $1 - f_1 > f_0/f$ is

$$x - 1 > x(1-\theta^{-1})^2 \tag{24}$$

There are two cases:

1. $\theta < 0.5$ in which case, the above inequality cannot be satisfied. In this case, as $\alpha$ increases, $\mu$ increases due to increase in $f_0$ until capacity is reached.

2. $\theta > 0.5$, in this case, for sufficiently small $\alpha$ such that

$$x > \frac{1}{2\theta^{-1} - \theta^{-2}} \tag{25}$$

$\mu$ decreases due to the increase in $1 - f_1$. Once

$$x < \frac{1}{2\theta^{-1} - \theta^{-2}} \tag{26}$$

$\mu$ increases due to increase in $f_0$ until capacity $x = 1$ is reached.

#### 2.3 Weak retrieval states

The fixed points $(m^*, \mu^*)$ corresponding to stable states of the network dynamics, are described by

$$\begin{aligned} m^* &= H\left(\frac{\theta-(1-f)m^*}{\sqrt{\alpha\mu^*}}\right) - H\left(\frac{\theta+fm^*}{\sqrt{\alpha\mu^*}}\right) \\ \mu^* &= fH\left(\frac{\theta-(1-f)m^*}{\sqrt{\alpha\mu^*}}\right) + (1-f)H\left(\frac{\theta+fm^*}{\sqrt{\alpha\mu^*}}\right) \end{aligned} \tag{27}$$

Numerical solutions of these equations reveal four qualitatively different kinds of fixed points. There exist two types of background states that do not overlap with the stored patterns ($m^* = 0$): one in which all the neurons in the network are silent $\mu^* = 0$ (*silent* state), and one where a finite fraction of the neurons are active, $\mu > f$ (*active background* state). In addition, there exist two types of retrieval states that have a finite overlap with one of the stored patterns ($m^* > 0$): one that overlaps almost perfectly with the pattern, $m^* \simeq 1, \mu^* \simeq f$ (*strong retrieval* state), and one in which there is a finite fraction of active background neurons, $m^* = O(1)$, $\mu^* > f$, and therefore the overlap with the corresponding pattern is weaker (*weak retrieval* state). Note that to our knowledge the active background state and the weak retrieval state had not been identified previously as solutions to these equations.

As mentioned above equations (27) have been studied previously in the limit of small $f$ [Tsodyks and Feigel'man, 1988, Sompolinsky and White, 2003]. In this limit the background states ($m^* = 0$) are the solution of

$$\mu^* = H\left(\frac{\theta}{\sqrt{\alpha\mu^*}}\right) \tag{28}$$

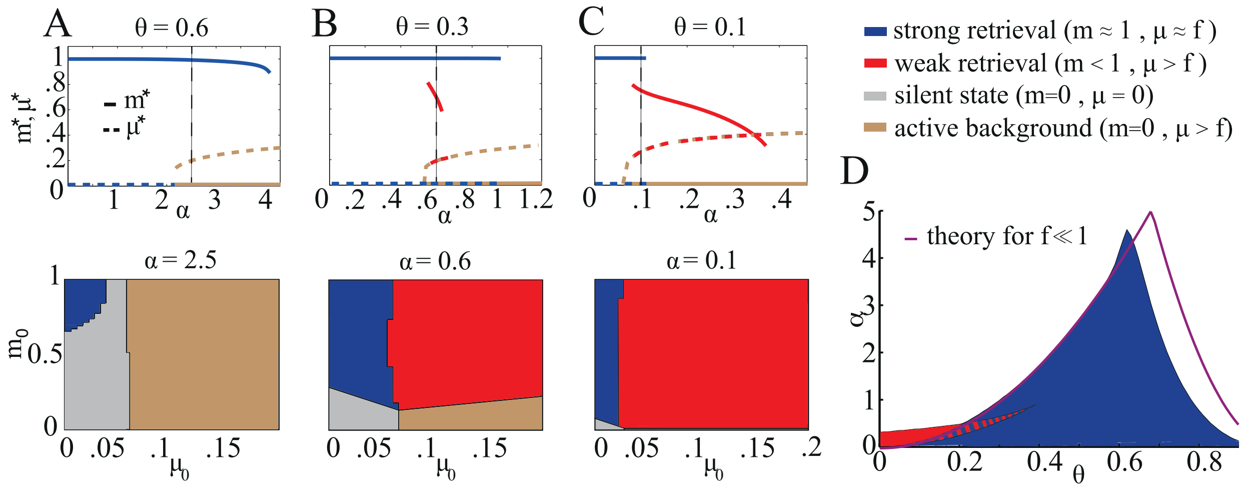


Fig S 1: *Fixed points* $(m^*, \mu^*)$ *of the map* $(m(t), \mu(t)) \rightarrow (m(t+1), \mu(t+1))$ *for patterns of coding level* $f = 0.01$ *at* $T = 0$*. A. Top panel, fixed points* $(m^*, \mu^*)$ *as a function of the memory load* $\alpha$ *for* $\theta = 0.6$*. Two kinds of fixed points are shown, the strong retrieval state (blue curves, full:* $m^*$*; dashed:* $\mu^*$*) and the active background state (brown curves). The silent state (not shown) is present for all* $\alpha$*, but is not shown here. Bottom panel shows which initial conditions* $(m_0, \mu_0)$ *lead to a given state, for* $\alpha = 2.5$ *and* $\theta = 0.6$*. B. Same as A but for* $\theta = 0.3$*. In addition to the previously mentioned fixed points, there is another retrieval state with high activity (weak retrieval state, red curves). C. Same as A and B, but for* $\theta = 0.1$*. D. Storage capacity of the network, i.e. the largest* $\alpha$ *for which there exists a retrieval state, as a function of* $\theta$*. For each value of* $\theta$ *the blue region correspond to the range of* $\alpha$ *for which the strong retrieval state exists. The red region corresponds to the existence of the weak retrieval state. The regions with blue and red stripes is where the two retrieval states are observed simultaneously.*

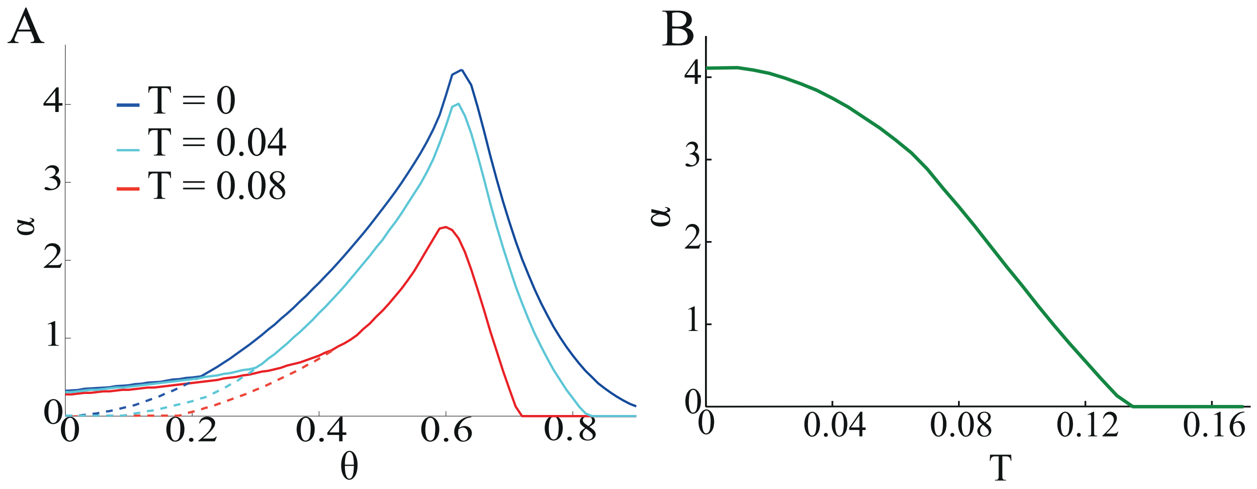


Fig S 2: *Effect of noise on memory properties of a single layer, for $f = 0.01$. A. Storage capacity as a function of $\theta$ for two values of the temperature (blue line is for $T = 0$ ; magenta line for $T = 0.08$). Above the dashed curves, only the weak retrieval state exists. B. Storage capacity as a function of temperature, for $\theta = 0.6$.*

For $\alpha < \alpha_{bg} < \alpha_c$ this equation has a single solution $\mu^* = 0$, while for $\alpha > \alpha_{bg}$, another stable solutions appears with $\mu^* \neq 0$, which we call the active background state.

The strong retrieval state has already been characterized in the main text. But when solving the fixed point equations numerically, we also found a retrieval state ($m^* = O(1)$) with a high value of $\mu^*$ (weak retrieval state). The region of existence of the weak retrieval state in the $\theta - \alpha$ plane is shown in red in figure Fig. S1A. It exists only for low activation threshold that are sub-optimal when considering performance of the TRN.

We have also studied these equations for different values of $f$, the coding level of memories. The trend we observed is that as $f$ increases the storage capacity gets smaller and is reached for smaller values of the activation threshold (e.g. for $f = 0.1$, $\alpha_c = 0.6$ at $\theta = 0.45$). Also, as $f$ increases from $0.01$ the size of the region where the two retrieval states coexist is reduced.

#### 2.4 Effects of temperature

We also quantified how storage capacity of the single layer network is decreased when noise is taken into account. For $f = 0.01$, figure S2A shows the storage capacity as a function of $\theta$ for $T = 0$ (blue line) $T = 0.04$ (cyan line) and $T = 0.08$ (magenta line). Note that the weak retrieval state is more robust to noise than the strong retrieval state. Figure S2B illustrates how storage capacity drops when noise is increased for $f = 0.01$ and $\theta = 0.6$. There is a critical temperature beyond which the storage capacity vanishes, as expected from previous studies of attractor neural networks [Amit, 1989].

### 3 Single path

#### 3.1 Dynamical equation for propagation of activity

When a sequence of patterns $\vec{\eta}(t)$, drawn independently from stored patterns, is fed to the first layer of the feed-forward chain, it elicits activity without eliciting pattern retrieval in layers. In layer $l$ this activity can be characterized by the macroscopic parameters:

$$n^l(t) = \frac{1}{Nf(1-f)} \sum_i^N \langle (\eta_i(t-l) - f)\sigma_i^l(t) \rangle \tag{29}$$

the overlap of the state of layer $l$ with $\vec{\eta}(t)$ of sparseness f. And the mean activity in layer $l$:

$$\mu^l(t) = \frac{1}{N} \sum_i^N \langle \sigma_i^l(t) \rangle \tag{30}$$

To express those macroscopic variables one time step later, one can proceed as in the previous section:

$$n^l(t+1) = f_1^l(t+1) - f_0^l(t+1) \tag{31}$$

where $f_1^l(t+1) = \langle \sigma_i^l(t+1) \rangle_{\eta_i^1=1}$ (*resp.*$f_0^l(t+1) = \langle \sigma_i^l(t+1) \rangle_{\eta_i^1=0}$) is the average of $\sigma_i^l(t+1)$ for sites i such that $\eta_i^1 = 1$ (*resp.* $\eta_i^1 = 0$).
Each of these average depends one whether sites are active or not in the previous layer, the time step before:

$$\begin{aligned} f_1^l(t+1) &= f_1^{l-1}(t) \int_{-\infty}^{+\infty} \frac{dz}{\sqrt{2\pi}} e^{-\frac{z^2}{2}} \frac{1}{1 + \exp(-\frac{1}{T}[\lambda\sqrt{\frac{p}{C}\mu^l(t)}z - \theta + 1])} \\ &\quad + (1 - f_1^{l-1}(t)) \int_{-\infty}^{+\infty} \frac{dz}{\sqrt{2\pi}} e^{-\frac{z^2}{2}} \frac{1}{1 + \exp(-\frac{1}{T}[\lambda\sqrt{\frac{p}{C}\mu^l(t)}z - \theta])} \end{aligned} \tag{32}$$

and

$$\begin{aligned} f_0^l(t+1) &= f_0^{l-1}(t) \int_{-\infty}^{+\infty} \frac{dz}{\sqrt{2\pi}} e^{-\frac{z^2}{2}} \frac{1}{1 + \exp(-\frac{1}{T}[\lambda\sqrt{\frac{p}{C}\mu^l(t)}z - \theta + 1])} \\ &\quad + (1 - f_0^{l-1}(t)) \int_{-\infty}^{+\infty} \frac{dz}{\sqrt{2\pi}} e^{-\frac{z^2}{2}} \frac{1}{1 + \exp(-\frac{1}{T}[\lambda\sqrt{\frac{p}{C}\mu^l(t)}z - \theta])} \end{aligned} \tag{33}$$

$f_1^l(t)$ and $f_0^l(t)$) are related to $n^l(t)$ and $\mu^l(t)$ by:

$$\begin{cases} n^l(t) &= f_1^l(t) - f_0^l(t) \\ \mu^l(t) &= f f_1^l(t) + (1-f) f_0^l(t) \end{cases} \Leftrightarrow \begin{cases} f_1^l(t) &= (1-f)n^l(t) + \mu^l(t) \\ f_0^l(t) &= -f n^l(t) + \mu^l(t) \end{cases} \tag{34}$$

Which gives:

$$
\begin{aligned}
n^l(t+1) &= n^{l-1}(t)\int_{-\infty}^{+\infty}\frac{dz}{\sqrt{2\pi}}e^{-\frac{z^2}{2}}\left[\frac{1}{1+\exp(-\frac{1}{T}[\lambda\sqrt{\frac{p}{C}\mu^l(t)}z-\theta+1])}\right. \\
&\left. -\frac{1}{1+\exp(-\frac{1}{T}[\lambda\sqrt{\frac{p}{C}\mu^l(t)}z-\theta])}\right] \qquad (35)
\end{aligned}
$$

And for the mean activity in layer $l$:

$$
\begin{aligned}
\mu^l(t+1) &= f\langle\sigma_i^l(t+1)\rangle_{\eta_i^1=1}+(1-f)\langle\sigma_i^l(t+1)\rangle_{\eta_i^1=0} \\
&= \mu^{l-1}(t)\int_{-\infty}^{+\infty}\frac{dz}{\sqrt{2\pi}}e^{-\frac{z^2}{2}}\Big(\frac{1}{1+\exp(-\frac{1}{T}[\lambda\sqrt{\frac{p}{C}\mu^l(t)}z-\theta+1])} \\
&-\frac{1}{1+\exp(-\frac{1}{T}[\lambda\sqrt{\frac{p}{C}\mu^l(t)}z-\theta])}\Big)\Big) \\
&+\int_{-\infty}^{+\infty}\frac{dz}{\sqrt{2\pi}}e^{-\frac{z^2}{2}}\frac{1}{1+\exp(-\frac{1}{T}[\lambda\sqrt{\frac{p}{C}\mu^l(t)}z-\theta])} \qquad (36)
\end{aligned}
$$

#### 3.2 The different regimes of mean activity

##### 3.2.1 Single layer:

To understand the behavior of the system, let us consider a single layer of neurons receiving input in the form of a sequence of random binary vectors $\eta_i(t)$ with a sparseness level $g$ and ask how the population's activity varies as $g$ increases from $g = f$. The equation describing this situation is given by

$$\mu(t+1) = gH\left(\frac{\theta-1}{\lambda\sqrt{\alpha\mu(t)}}\right)+(1-g)H\left(\frac{\theta}{\lambda\sqrt{\alpha\mu(t)}}\right) \qquad (37)$$

The fixed point equation is

$$\mu = gH\left(\frac{\theta-1}{\lambda\sqrt{\alpha\mu}}\right)+(1-g)H\left(\frac{\theta}{\lambda\sqrt{\alpha\mu}}\right) \qquad (38)$$

From that analysis of this equation, similarly to the capacity analysis of a single module, we expect that for a fixed $\alpha$ there are two regimes of $\mu$: for low $g$, $\mu$ is approximately $g$; above a critical $g$, $\mu$ is of order 1. To determine the transition, we write

$$g = \frac{1}{s}f \qquad (39)$$

where large values of $s$ means small $g$. Further, we denote

$$y = \frac{\theta^2}{2\lambda^2\alpha g|\log g|}, y > 1 \qquad (40)$$

implying,

$$\mu \approx g\left(1 + g^{y-1} - g^{y(1-\theta^{-1})^2}\right) \tag{41}$$

To relate $y$ to $s$, we use,

$$\alpha = \frac{\theta^2}{2\lambda^2 x f |\log f|}, x > 1 \tag{42}$$

yielding,

$$y = x\frac{f|\log f|}{g|\log g|} \approx sx \tag{43}$$

Thus, below the transition, we have

$$\mu \approx g\left(1 + g^{sx-1} - g^{sx(1-\theta^{-1})^2}\right) \tag{44}$$

and the transition occurs when $y = 1$ or $s = 1/x$, i.e.,

$$g_C = xf \tag{45}$$

For larger $g$, $\mu$ increases rapidly to order 1. How does $\mu$ behaves as a function of $g$ below the transition? As before, there are multiple regimes.

**Regime Ia:** $\theta < 0.5$
In which case, $\mu$ is monotonically increasing with $g$ until $sx = 1$ forcing a transition to large $\mu$.

**Regime Ib**: $\theta > 0.5$ and $x > \frac{1}{2\theta^{-1}-\theta^{-2}}$
In this case, initially ($s = 1$) and

$$sx > \frac{1}{2\theta^{-1} - \theta^{-2}} \tag{46}$$

hence $\mu$ decreases and $s$ increases, because the term $g^{sx(1-\theta^{-1})^2}$ is larger than $g^{sx-1}$. Increasing $s$ ensures that (46) holds hence, $\mu$ continues to decrease.

**Regime II:** $\theta > 0.5$ and $x < \frac{1}{2\theta^{-1}-\theta^{-2}}$
Here intially $sx = x$ and

$$sx < \frac{1}{2\theta^{-1} - \theta^{-2}} \tag{47}$$

hence $\mu$ increases until $sx = 1$ triggering a transition to large $\mu$.

##### 3.2.2 Propagation across layers:

We now turn to the multi-layer equations, and consider the equilibrium profile of activities, $\mu^l = \mu^l(\infty)$, given by

$$\mu^l = \mu^{l-1}\left[H\left(\frac{\theta-1}{\lambda\sqrt{\alpha\mu^l}}\right) - H\left(\frac{\theta}{\lambda\sqrt{\alpha\mu^l}}\right)\right] + H\left(\frac{\theta}{\lambda\sqrt{\alpha\mu^l}}\right) \tag{48}$$

with the boundary condition $\mu^1 = f$. To study the solution, we approximate the equations for $\mu$ using the scaling,

$$\alpha = \frac{\theta^2}{2\lambda^2 x f|\log f|}, \tag{49}$$

$$\frac{\mu^{-1}\theta^2}{2\lambda^2\alpha} = \mu^{-1} x f|\log f| \tag{50}$$

and denote, $\mu^{-1} = sf^{-1}$. Thus, $\frac{\mu^{-1}\theta^2}{2\lambda^2\alpha} = sx|\log f|$ and

$$H\left(\frac{\theta}{\lambda\sqrt{\alpha\mu}}\right) \approx \frac{\lambda\sqrt{\alpha\mu}}{\theta\sqrt{2\pi}}\exp\left(\frac{-\theta^2}{2\lambda^2\alpha\mu}\right) \approx \frac{1}{2\sqrt{\pi}\sqrt{sx|\log f|}} f^{sx} \tag{51}$$

$$H\left(\frac{1-\theta}{\lambda\sqrt{\alpha\mu}}\right) \approx \frac{\lambda\sqrt{\alpha\mu}}{(1-\theta)\sqrt{2\pi}}\exp\left(\frac{-(1-\theta)^2}{2\lambda^2\alpha\mu}\right) \approx \frac{1}{2\sqrt{\pi}(\theta^{-1}-1)\sqrt{sx|\log f|}} f^{sx(1-\theta^{-1})^2} \tag{52}$$

**Regime Ia**: $\theta < 0.5$.
According to the single layer analysis, we expect here that $\mu^l$ increases monotonically with $l$. The activity grows initially from an initial value of $\mu = f$ until it reaches the value $\mu = xf$ (corresponding to the critical $g$ in the single layer) at a critical $l$. Beyond that, there is a rapid increase of $\mu$ to a value close to 1. To evaluate the critical propagation length, we note that

$$H\left(\frac{\theta}{\lambda\sqrt{\alpha\mu^l}}\right) > H\left(\frac{1-\theta}{\lambda\sqrt{\alpha\mu^l}}\right) \tag{53}$$

Hence, we can approximate equation of $\mu$

$$\mu^l = \mu^{l-1}\left[1 - H\left(\frac{\theta}{\lambda\sqrt{\alpha\mu^l}}\right) - H\left(\frac{1-\theta}{\lambda\sqrt{\alpha\mu^l}}\right)\right] + H\left(\frac{\theta}{\lambda\sqrt{\alpha\mu^l}}\right) \tag{54}$$

by,

$$\mu^l = H\left(\frac{\theta}{\lambda\sqrt{\alpha\mu^l}}\right) + \mu^{l-1}\left[1 - H\left(\frac{\theta}{\lambda\sqrt{\alpha\mu^l}}\right)\right] \tag{55}$$

$$d\mu^l = \mu^l - \mu^{l-1} = H\left(\frac{\theta}{\lambda\sqrt{\alpha\mu^l}}\right) \tag{56}$$

$$d\mu^l \approx \frac{dl}{2\sqrt{\pi}\sqrt{sx|\log f|}} f^{sx} \tag{57}$$

$$ds = -\frac{1}{2f}s^2 d\mu = -\frac{dl}{2\sqrt{\pi}\sqrt{sx|\log f|}} s^2 f^{sx-1} \tag{58}$$

This allows to express $l$ as a function of $s$ and thus $L_c = l(s = 1/x)$, hence equation (**??**).

**Regime Ib**: $\theta > 0.5$ and $x < \frac{1}{2\theta^{-1}-\theta^{-2}}$
In this case, an approximate equation for $\mu^l$ is:

$$\mu^l = \mu^{l-1}\left[1 - H\left(\frac{1-\theta}{\lambda\sqrt{\alpha\mu^l}}\right)\right] + H\left(\frac{\theta}{\lambda\sqrt{\alpha\mu^l}}\right) \tag{59}$$

or,

$$d\mu^l = -\mu^l H\left(\frac{1-\theta}{\lambda\sqrt{\alpha\mu^l}}\right) + H\left(\frac{\theta}{\lambda\sqrt{\alpha\mu^l}}\right) \tag{60}$$

hence,

$$ds = \frac{dl}{2\sqrt{\pi}(\theta^{-1}-1)\sqrt{sx|\log f|}} s f^{sx(1-\theta^{-1})^2} - \frac{dl}{2\sqrt{\pi}\sqrt{sx|\log f|}} s^2 f^{sx-1} \tag{61}$$

and

$$l(s) = 2\sqrt{\pi}\int_s^1 \left[\frac{1}{\sqrt{sx|\log f|}} s^2 f^{sx-1} - \frac{1}{(\theta^{-1}-1)\sqrt{sx|\log f|}} s f^{sx(1-\theta^{-1})^2}\right]^{-1} ds \tag{62}$$

Once again, the critical length is given by the equation, $L_C = l(1/x)$, although it is hard to give a simple expression for $L_C$. To summarize, we can conclude from (62), that the typical decay length, $l_c$ of $\mu$ in this regime is given as

$$1/l_c \propto \frac{1}{(\theta^{-1}-1)\sqrt{x|\log f|}} f^{x(1-\theta^{-1})^2} - \frac{1}{\sqrt{x|\log f|}} f^{x-1} \tag{63}$$

**Regime II**: $\theta > 0.5$ and $x > \frac{1}{2\theta^{-1}-\theta^{-2}}$ Here the propagation of the random sequence does not 'collapse' at a finite critical length. Instead, $\mu$ decreases with $l$ smoothly. To estimate the characteristic decay length,
We observe that due to the constraint on $x$ relative to this regime, $ds$ has a positive sign, implying that $\mu$ decreases with $l$. Hence, we have here,

$$l(s) = 2\sqrt{\pi}\int_1^s \left[\frac{1}{(\theta^{-1}-1)\sqrt{sx|\log f|}} s f^{sx(1-\theta^{-1})^2} - \frac{1}{\sqrt{sx|\log f|}} s^2 f^{sx-1}\right]^{-1} ds \tag{64}$$

where $s(l = 0) = 1$, $s(l) \geq 1$.

##### 3.3 Dynamical equations for memory retrieval

If pattern $\vec{\xi}^{1,l_0}$ is sent to the first layer, it will elicit pattern retrieval in $l_0$. We are interested in calculating the time evolution of

$$m^{l_0}(t) = \frac{1}{Nf(1-f)} \sum_i^N \langle (\xi_i^{1,l_0} - f)\sigma_i^{l_0}(t) \rangle \tag{65}$$

The field on a neuron $i$ is given by

$$\begin{aligned} h_i^{l_0}(t) &= \sigma_i^{l_0-1}(t) + \lambda \frac{1}{Cf(1-f)} (\xi_i^{1,l_0} - f) \sum_{r=1}^K (\xi_{j_r}^{1,l_0} - f)\sigma_{j_r}(t) \\ &+ \lambda \frac{1}{Cf(1-f)} \sum_{\mu=2}^p (\xi_i^{\mu,l_0} - f) \sum_{r=1}^K (\xi_{j_r}^{\mu,l_0} - f)\sigma_{j_r}(t) \end{aligned} \tag{66}$$

Once again, the overlap one time step after can be written

$$m^{l_0}(t+1) = \langle \sigma_i^{l_0}(t+1) \rangle_{\xi_i^{1,l_0}=1} - \langle \sigma_i^{l_0}(t+1) \rangle_{\xi_i^{1,l_0}=0} \tag{67}$$

In this case, those averages are given by

$$\langle \sigma_i^{l_0}(t+1) \rangle_{\xi_i^{1,l_0}=1} = f_1^{l_0}(t+1) = f_1^{l_0-1}(t) I(1,1) + (1 - f_1^{l_0-1}(t)) I(1,0) \tag{68}$$

where we have introduced the notation:

$$I(x,y) = \int_{-\infty}^{+\infty} \frac{dz}{\sqrt{2\pi}} e^{-\frac{z^2}{2}} \frac{1}{1 + \exp(-\frac{1}{T}[\lambda(x-f)m^{l_0}(t) + \lambda\sqrt{\alpha\mu^{l_0}(t)}z - \theta + y])}. \tag{69}$$

Note that I(x,y) depends on $m^{l_0}(t)$ and $\mu^{l_0}(t)$, and that $f_1^{l_0}(t)$ (resp. $f_0^{l_0}(t)$) is the probability that a neuron $i$ of layer $l_0$ such that $\xi_i^{l_0} = 1$ (resp. $\xi_i^{l_0} = 0$) is active.

$$\langle \sigma_i^{l_0}(t+1) \rangle_{\xi_i^{l_0}=1} = f_0^{l_0}(t+1) = f_0^{l_0-1}(t) I(0,1) + (1 - f_0^{l-1}(t)) I(0,0) \tag{70}$$

Using the relationships (34) between $f_1^{l_0-1}(t)$, $f_0^{l_0-1}(t)$, $m^{l_0}(t)$ and $\mu^{l_0}(t)$, we have

$$\begin{aligned} m^{l_0}(t+1) &= m^{l_0-1}(t)\{(1-f)[I(1,1) - I(1,0)] + \\ &\quad +f[I(0,1) - I(0,0)]\} \\ &\quad +\mu^{l_0-1}(t)\{I(1,1) - I(1,0) \\ &\quad -[I(0,1) - I(0,0)]\} \\ &\quad +I(1,0) - I(0,0) \end{aligned} \tag{71}$$

The recurrence relationship for the mean activity is

$$\begin{aligned} \mu^{l_0}(t+1) &= f\langle \sigma_i^{l_0}(t+1) \rangle_{\xi_i^{l_0}=1} + (1-f)\langle \sigma_i^{l_0}(t+1) \rangle_{\xi_i^{l_0}=0} \\ &= m^{l_0-1}(t) f(1-f)\{I(1,1) - I(1,0) \\ &\quad -[I(0,1) - I(0,0)]\} \\ &\quad +\mu^{l_0-1}(t)\{f[I(1,1) - I(1,0)] + \\ &\quad +(1-f)[I(0,1) - I(0,0)]\} \\ &\quad +fI(1,0) + (1-f)I(0,0) \end{aligned} \tag{72}$$

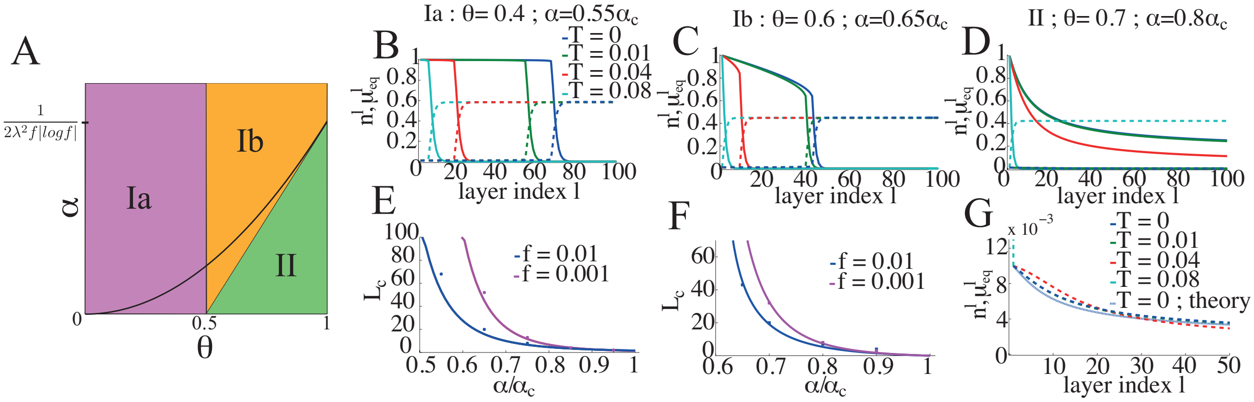


Fig S 3: *Propagation of inputs and average activity in steady state. A. Regions in the $\theta - \alpha$ plane where the different regimes are found in the case $T = 0$ and $f \ll 1$. The parabola in black represents the storage capacity $\alpha_c$. B. Dashed lines represent $\mu^l_{eq}$ the average activity at equilibrium in each layer in regime Ia. Full lines represent the overlap $n^l$ between a pattern sent at time $t$ and the state of layer $l$ at time $t+l$. The coding level of inputs is $f = 0.01$. C-D. Same as A for regime Ib and II. E. $L_c$ as a function of the normalized storage load $\frac{\alpha}{\alpha_c}$ in regime Ia. Full lines correspond to the analytical expression for $f \ll 1$, and dots are obtained by integrating the dynamical equations and measuring the layer for which a sharp increase in average activity occurs. F. Same as D in regime Ib. G. Zoom on the average activity in regime II. Different colors correspond to different temperatures as in panels A-C.*

#### 3.4 Effect of $\lambda$ on storage capacity at $T = 0$

Introducing the parameter $\lambda$ scaling the recurrent inputs compared to feed-forward ones as little effects on the capacity of a path as the optimal value is close to 1 as shown in figure S4 where color coded is $\alpha_T$ as a function of $\theta$ and $\lambda$ for a path of depth $L = 1,000$.

#### 3.5 Storage capacity of a path at T>0

The performance of the CRN is measured by $\alpha_T$ which is proportional to the total number of patterns that can be retrieved and maintained by cueing with the root module,

$$\alpha_T = \alpha L_r \tag{73}$$

where $\alpha$ is the storage load of each layer, and $L_r(\alpha)$ is the deepest layer for which pattern retrieval is possible. In qualitative terms, in order to have a large value of $\alpha_T$ there should be as many patterns as possible stored in each layer, while propagation of patterns through the layers of the CRN should be as faithful as possible in order to allow retrieval in deep layers (large value of $L_r(\alpha)$).

**Computation of $\alpha_T$**

It is difficult to obtain an analytical formula for $L_r(\alpha)$ because of the difficulty to get a formula for how the overlap $n^l$ decreases with $l$. To have quantitative estimates of $\alpha_T$, we iterate the dynamical equations for the order parameters. Equations are first iterated to set the mean activities at $\mu^l_{eq}$, then a pattern $\vec{\xi}^{\mu,l_0}$ is presented to the first layer, and its

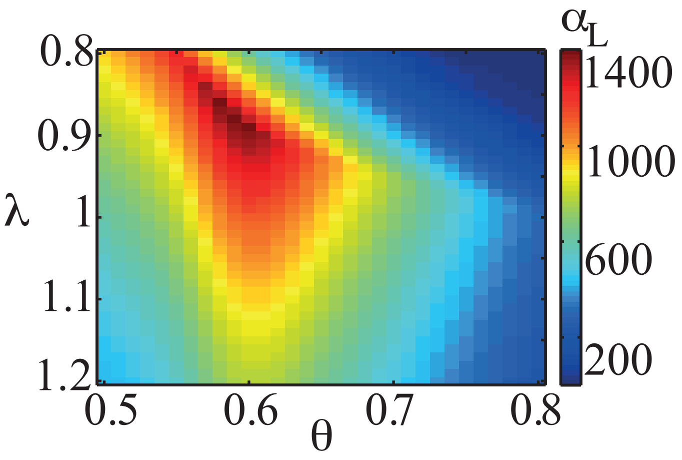


Fig S 4: *Dependence of storage capacity of a path on* $\lambda$ *for* $L = 1,000$

propagation is evaluated by iterating (35),(36) until layer $l_0$ is reached, then the evolution of $m^{l_0}$ is monitored by iterating (71),(72). Retrieval is successful if $m^{l_0}(t \gg l_0) \simeq 1$, and, for a given value of the storage load in each layer $\alpha$, $L_r(\alpha)$ can be measured by looking for the deepest layer for which retrieval is possible. Thus $\alpha_T$ can be evaluated.

**Storage capacity at** $T > 0$

The same principles hold for non-zero temperatures. Figure S5C shows $\alpha_T$ as a function of $\alpha$ for $\lambda = 1$, $T = 0.08$, $L = 100$ for different values of $\theta$. At non-zero temperatures, the initial increase with $\alpha$ is not linear anymore because already for small $\alpha$, patterns in the deepest layers can not be retrieved by cueing the first layer. For these parameters, the optimal capacity is reached at $\alpha = 0.5 \simeq \frac{\alpha_c}{3}$, and patterns can be retrieved in 22 layers as can be seen in figure S5E. The maximal capacity is reached for $\theta \simeq 0.6$. Figures S5D and F are similar to C and E, for a fixed value $\theta = 0.6$ and different values of $\lambda$. Optimal capacities are reached for $\lambda \simeq 1.1 - 1.2$.

#### 3.6 Reading-out in the last module of a path

When a pattern $\vec{\xi}^{\mu,l_0}$ is sustained in layer $l_0$, the inputs sent to downstream layers ($l > l_0$) represent both $\vec{\xi}^{\mu,l_0}$ and the flow of inputs $\vec{\eta}(t)$ that are presented to the first layer of the CRN. The quality of the representation of $\vec{\xi}^{\mu,l_0}$ in these downstream layers depends dramatically on the profile of average activities $\mu^{l>l_0}$. It behaves as the profile of average activity obtained in previous sections with a different boundary between regime Ib and regime II given by $\alpha = \frac{1}{1+\mu^{l_0}/f}\frac{\theta-\frac{1}{2}}{\lambda^2 f|\ln f|}$. In regime Ia-b when $\mu^l$ increases with $l$, the quality of the representation of $\vec{\xi}^{\mu,l_0}$ decreases slowly as $l$ is increased until a layer $L_c$ for which average activity increases sharply and becomes of order one. This is illustrated in figure S6A where we show the overlap $m^{l,l_0}$ between $\vec{\xi}^{\mu,l_0}$ and $\vec{\sigma}^l$ in regime Ia for different values of $\alpha$. In figure S6B we show the same quantities for $\theta = 0.6$ with different values of $\alpha$ that samples both regime Ib (cyan and green lines) and II (blue and magenta lines). We also performed the same study for $T = 0.08$, $\lambda = 1$ and $f = 0.01$. Results are shown in figure

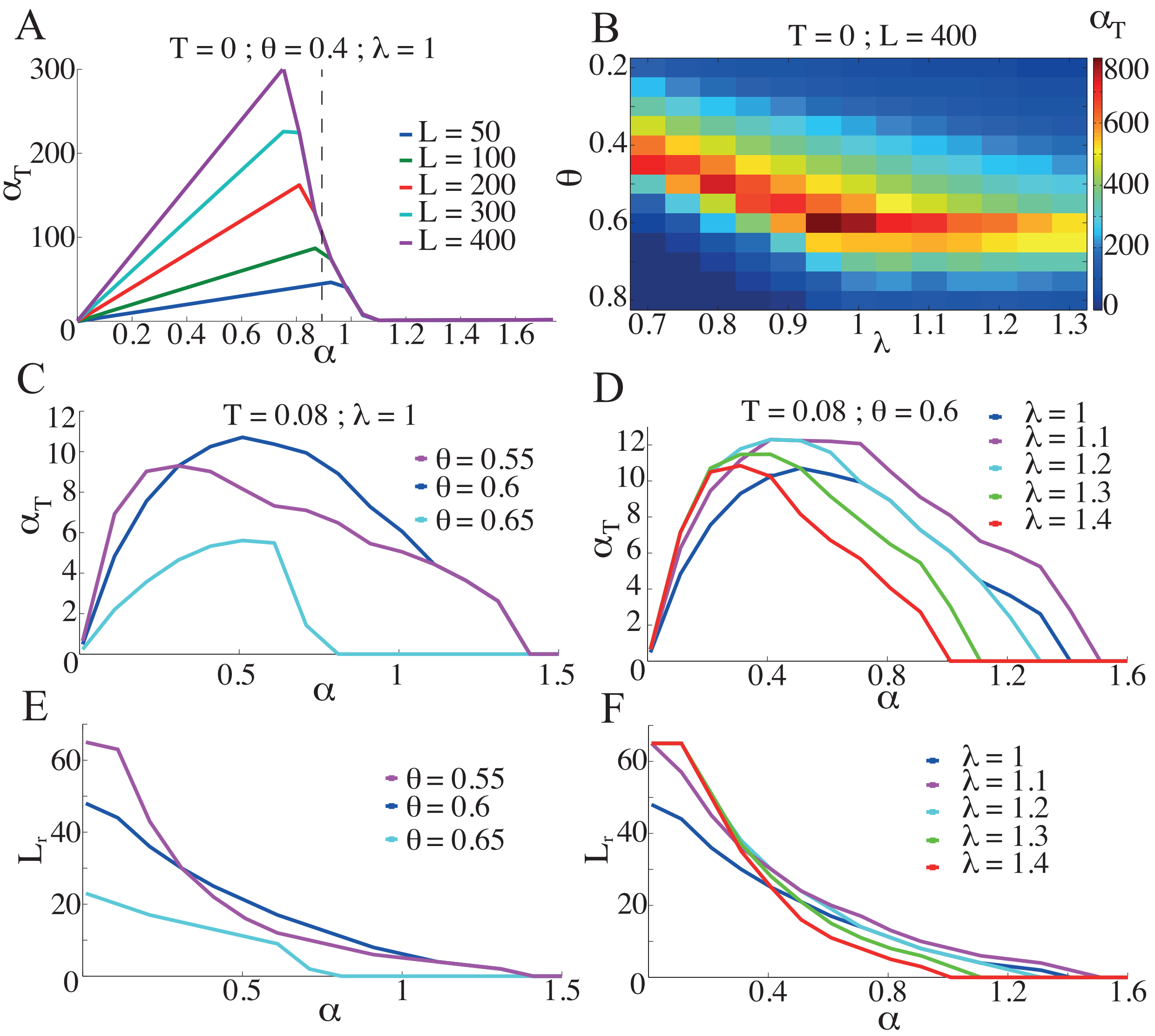


Fig S 5: *Storage capacity of the CRN, i.e. the maximal number of patterns that can be stored and retrieved by cueing the first layer, as defined in equation* (73). *A. Storage capacity for $T = 0$ and $f = 0.01$ as a function of $\alpha$ which quantifies the number of patterns stored in each layer. The dashed vertical line is at $\alpha = \frac{\alpha_c}{2}$ B. Storage capacity, optimized over $\alpha$, as a function of parameters $\theta$ and $\lambda$. C. Storage capacity as a function of $\alpha$ for $T = 0.08$ and $f = 0.01$ for different values of $\theta$. D. Same as C for different values of $\lambda$. E-F Values of $L_r$, the last layer of the CRN for which retrieval can be achieved by cueing the first layer, corresponding to the storage capacities obtained in panels C and D.*

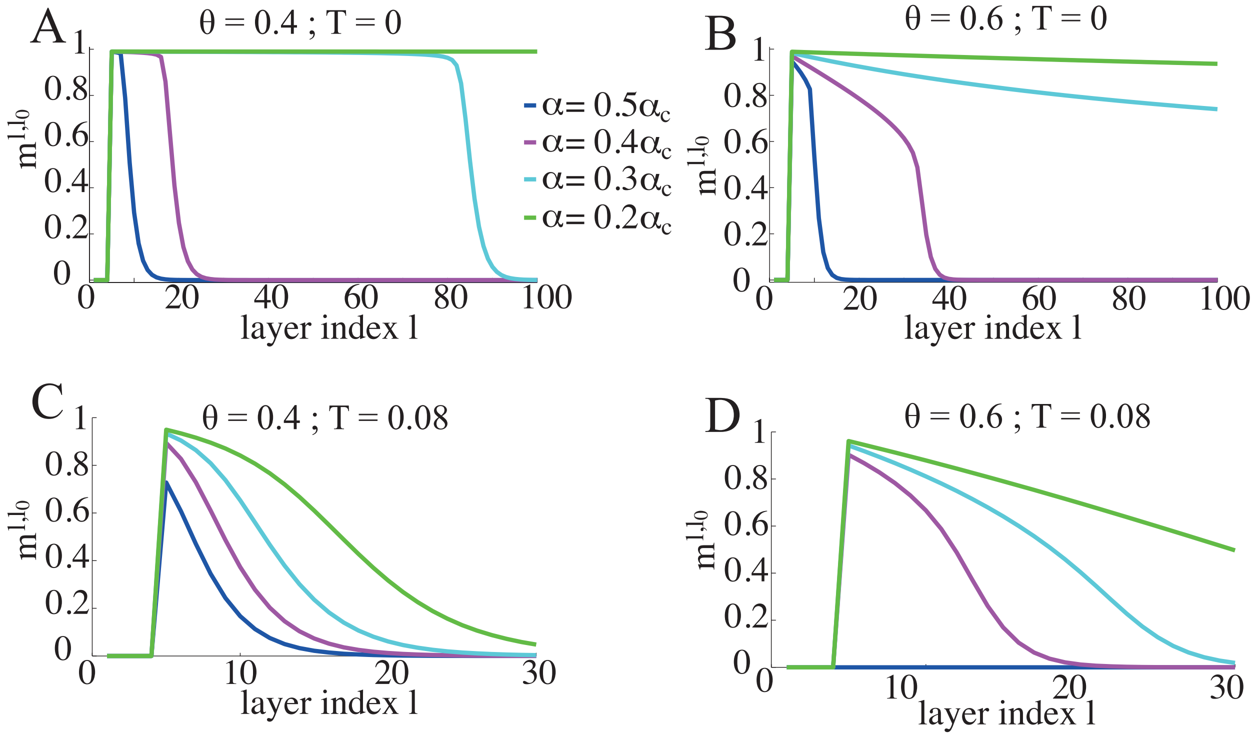


Fig S 6: *Read-out of a retrieved patterns by downstream layers. A pattern* $\vec{\xi}^{\mu_0,5}$ *is retrieved in layer 5 for a CRN with* $T =$ *and* $f = 0.01$*. We measure the overlap* $m^l$ *between* $\vec{\xi}^{\mu_0,5}$ *and* $\vec{\sigma}^{l>5}$ *to estimate the quality of the representation of the retrieved memory in downstream layers. A. Behavior of* $m^l$ *for* $\theta = 0.4$*. Behavior of* $m^l$ *for* $\theta = 0.6$*. C-D* $m^l$ *at* $T = 0.08$ *for* $\theta = 0.4$ *and* $\theta = 0.6$*.*

S6C and D. For a retrieval in layer 5, the retrieved pattern can be read-out by $\simeq 10$ layers, a number of the same order as $L_r$ the number of layers for which retrieval is possible (see figure S5E-F).

### 4 Numerical simulations

All the results we have presented so far have been derived for diluted networks in the large $N$ limit ($N \gg C \to \infty$). In this section we present the result of simulations of finite size networks that are fully connected ($C = N$). We first estimate the storage capacity of single layer networks of sizes $N = 1000$, $N = 5000$, $N = 10000$ for an activation threshold $\theta = 0.6$. In figure S7A we plot the two order parameters $m$ and $\mu$ once the network has reached a steady state, after it has been initialized in one of its memory state. This is done for two values of the noise.

At $T = 0$, networks of all three sizes have memory states for values of the memory load comparable to the maximal capacity predicted by the calculations in the $N \to \infty$. The transition at which the strong retrieval states disappear is not as sharp as predicted by the calculation. Above the maximal capacity, the system exhibits remanent overlaps with the memory state with which the system was initialized, similar to remanent magnetization in spin-glass models and the Hopfield model [Amit et al., 1987]. For instance, for $\alpha = 10\alpha_c$ and $N = 1,000$, we have $m^* \simeq 0.6$. Note that the state reached by the network has a large average activity ($\mu \simeq 0.15$) and in fact overlaps with multiple memories. At $T = 0.08$ , the network of size $N = 1000$ can not be used as a memory device, while larger networks have a storage capacity which is close to the analytical prediction.

We now turn to the full CRN, and validate some of the key results by simulating networks with $L = 25$ layers of $N = 5,000$ neurons, with the standard parameters ($T = 0.08$, $\theta = 0.6$, $\lambda = 1$). Figure S7B shows how a pattern propagates through the different layers. We present a pattern of activity (that does not correspond to any stored memory) to the first layer and measure how well it is represented in the different layers by computing the overlap $n^l$. Data shown in this figure are an average over 15 realizations.

We then test the retrieval ability of a CRN. Figure S7C shows the deepest layer for which a pattern can be retrieved (average over 15 realizations). For finite size networks, there is no sharp transition with the memory load $\alpha$. $L_c$ is now defined as the last layer for which more than half of the tested patterns are retrieved (here a pattern is said to be retrieved if after 50 time steps $m > 0.7$.) Despite some quantitative differences for large values of $\alpha$, these simulations suggest the result presented in the above sections can be extended to finite-size networks ($N \geq 5000$) with dense connectivity.

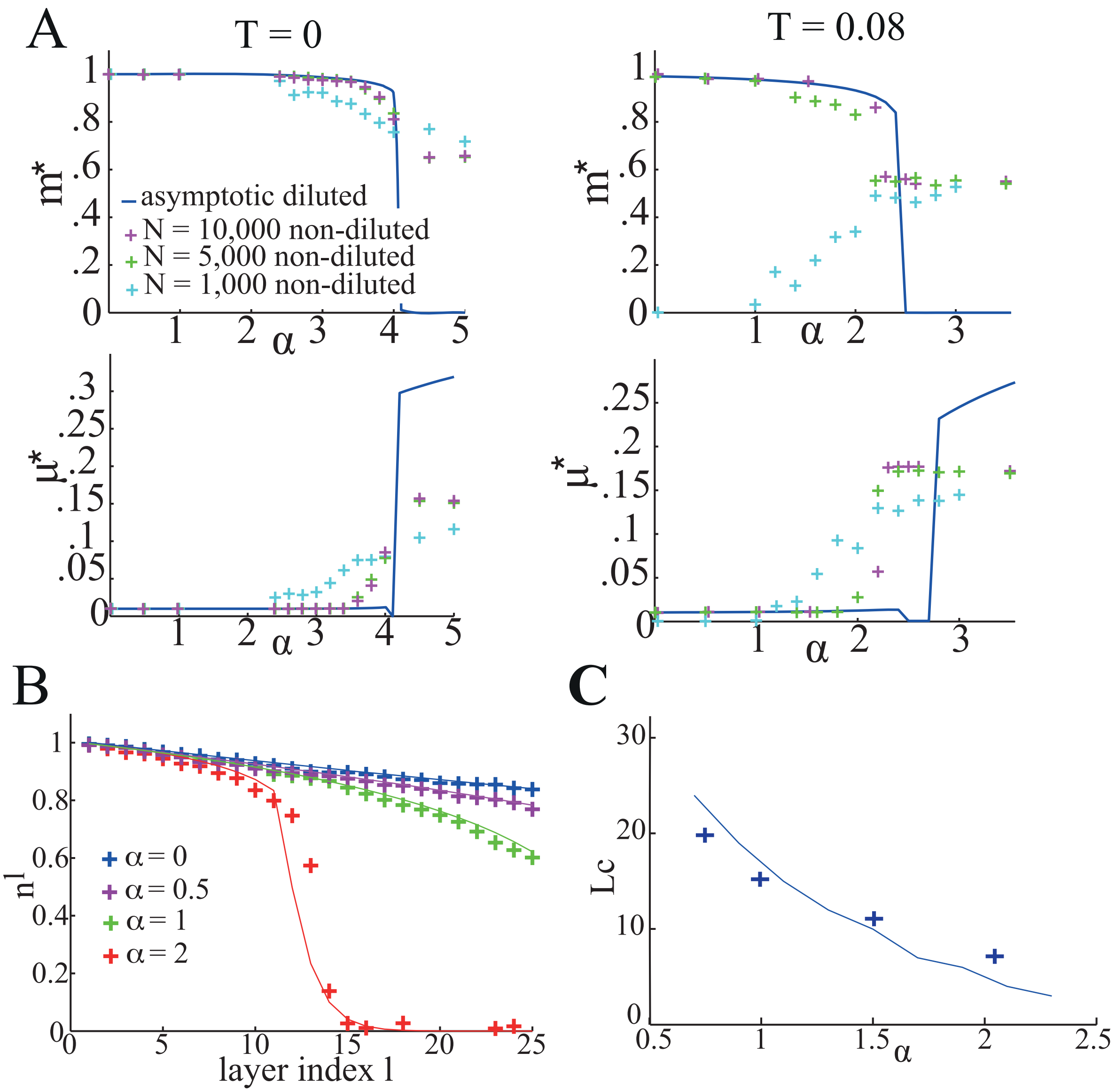


Fig S 7: *Numerical simulations. In all the panels solid lines correspond to analytical results for an extremely diluted network with* $N \to +\infty$, $\theta = 0.6$, $\lambda = 1$. *A. Single layer for various network sizes for a noise* $T = 0$ *(left panels) and* $T = 0.08$ *(right panels). The network is initialized in one of its memory state, the dynamics is iterated and the final state is characterized by the overlap with the initial pattern* $m^*$ *(up) and the average activity* $\mu^*$ *(bottom). Each cross corresponds to testing* 50 *patterns. Solid lines are for a diluted infinite size network,* $\theta = 0.6$, *and* $T = 0.08$ *here and in the other panels B. A pattern uncorrelated with the stored memories is sent to the first layer of a path (*$L = 25$, $N = 5000$, $\lambda = 1$*), and its representation in the subsequent layers as it travels through the network is measured by the overlap* $n^l$. *Each cross is an average of* $n^l$ *over* 15 *different patterns. C. Measure of* $L_c$ *the deepest layer in which a memory can be retrieved (*$L = 25$, $N = 5000$, $\lambda = 1$, $\theta = 0.6$*). In simulations* $L_c$ *is defined as the deepest layer for which more than half of* 15 *tested patterns are retrieved after they have been presented to the first layer.*

[Derrida et al., 1987] Derrida, B., Gardner, E., and Zippelius, A. (1987). An exactly solvable asymmetric neural network model. *Europhys. Lett.*, 4:167–173.

[Evans, 1989] Evans, M. (1989). Random dilution in a neural network for biased patterns. *Journal of Physics A: Mathematical and General*, 22(12):2103.

[Sejnowski, 1977] Sejnowski, T. J. (1977). Storing covariance with nonlinearly interacting neurons. *J. Math. Biol.*, 4:303–.

[Sompolinsky and White, 2003] Sompolinsky, H. and White, O. (2003). Theory of large recurrent networks: from spikes to behavior. In *Methods and Models in Neurophysics, Volume Session LXXX: Lecture Notes of the Les Houches Summer School*, page Chapter 8.

[Tsodyks and Feigel'man, 1988] Tsodyks, M. and Feigel'man, M. V. (1988). The enhanced storage capacity in neural networks with low activity level. *Europhys. Lett.*, 6:101–105.
